# Laminin N-terminus α31 regulates keratinocyte adhesion and migration through modifying the organisation and proteolytic processing of laminin 332

**DOI:** 10.1101/617597

**Authors:** Lee D. Troughton, Valentina Iorio, Liam Shaw, Conor J Sugden, Kazuhiro Yamamoto, Kevin J. Hamill

**Author notes:** These authors contributed equally to this work (Lee D. Troughton, Valentina Iorio). **Corresponding author:**, Department of Eye and Vision Science, Institute of Life Course and Medical Sciences, William Henry Duncan Building, 6 West Derby Street, University of Liverpool, UK, L7 8TX.

## Abstract

Laminin N-terminus α31 (LaNt α31), a member of the laminin superfamily, expressed at low levels in intact epithelium but upregulated during wound repair. Increased expression of LaNt α31 reduced migration rate of corneal keratinocytes through an unknown mechanism. Here, we investigated whether LaNt α31 influences cell behaviour through modulating laminin-mediated processes. Adenoviral delivery of LaNt α31 into corneal epithelial cells led to reduced migration speed and increased cell spreading and changed laminin 332 organisation from diffuse arcs to tight clusters. Enhanced recruitment of collagen XVII and bullous pemphigoid antigen 1e to β4 integrin, indicating early maturation of hemidesmosomes, and changed focal adhesion distribution were also identified. LaNt α31 and laminin β3 co-immunoprecipitated from doubly transduced cells and were deposited together in live imaging experiment. Moreover, LaNt α31 expression led to increased matrix metalloproteinase (MMP) activity and proteolytic processing of laminin α3, and the inhibition of MMP activity rescued the laminin and hemidesmosome phenotypes. Provision of cell-derived extracellular matrix rescued the cell spreading and motility effects. These findings reveal LaNt α31 as a new player in regulating cell-to-matrix adhesion through its ability to influence laminin organisation and proteolytic processing.

## Introduction

The laminins (LMs) are an essential family of structural proteins that are incorporated into specialised regions of extracellular matrix (ECM) termed basement membranes (BMs), which in intact tissue, provide structural support for sheets of epithelial or endothelial cells, as well as support points for nerves and muscle(Hamill *et al.*, 2009b; Hohenester and Yurchenco, 2013). LMs also provide an essential substrate for cell migration during tissue remodelling and wound repair. Transitioning between these related but opposing roles requires context-specific changes to the substrate presented to cells. Understanding the mechanisms driving these transitions could provide a route toward therapeutic intervention, particularly in cases where wounds either heal slowly or heal ineffectively such as in corneal recurrent erosion syndrome(Pal-Ghosh *et al.*, 2008; Pal-Ghosh *et al.*, 2016). Here we investigated a relatively unstudied member of the LM superfamily, termed Laminin N-terminus α31 (LaNt α31), that could be an important contributor to these processes.

LaNt α31 is widely expressed across multiple tissue types(Hamill *et al.*, 2009c; Troughton *et al.*, 2020a). In the ocular anterior surface epithelium, it is highly abundant in the limbal epithelium with comparatively lower levels found in corneal and conjunctiva in intact tissue. LaNt α31 expression also has been shown to increase in the cornea during the active phases of wound healing the and to normal levels once the wound has healed(Barrera *et al.*, 2018). Upregulation of LaNt α31 expression has also been identified in *ex vivo* models of limbal stem cell activation and been associated with breast cancer(Barrera *et al.*, 2018; Troughton *et al.*, 2020b). *In vitro* functional studies have revealed that knockdown in epidermal keratinocytes reduced cell adhesion strength and slowed the rate of scratch wound closure(Hamill *et al.*, 2009c), while overexpression in primary limbal keratinocytes also caused a reduction in migration rates (Barrera *et al.*, 2018). However, the mechanism behind these effects remains unknown.

LaNt α31 is produced by alternative splicing of the LM α3 encoding gene, LAMA3, *via* a process involving splice site read through of exon nine and intronic polyadenylation(Hamill *et al.*, 2009c). The resulting transcript encodes a protein which major structural features are a LM N-terminal domain (LN domain) followed by a short stretch of LM-type epidermal growth factor-like repeats (LE domains) and a short unique carboxyl-terminus with no homology to conserved-domains (Figure 1A). Importantly, whereas LMs are obligate heterotrimers comprised of one α chain, one β chain and one γ chain that assemble *via* a LM coiled-coil domain (LCC domain) toward the carboxy-terminus of each subunit(Hamill *et al.*, 2009b; Aumailley, 2013; Hohenester and Yurchenco, 2013), LaNt α31 does not contain this LCC domain and therefore exists as an independent LM short-arm fragment (Figure 1A)(Hamill *et al.*, 2009c); however, the presence of the LN domain does suggest the potential to interact with LMs. Specifically, LM higher-order network assembly involves interaction between LN domains(Yurchenco *et al.*, 1985; Hussain *et al.*, 2011; Purvis and Hohenester, 2012; Hohenester and Yurchenco, 2013). Moreover, a LM-superfamily member, netrin-4, which consists of an LN domain with partial homology to LM β-chain LN domains, is capable of preventing LM-LM interactions and disrupting preformed networks(Schneiders *et al.*, 2007; Reuten *et al.*, 2016). Consistent with this concept, recent findings have revealed that induced LaNt α31 expression in breast cancer cells modifies their invasion characteristics into a LM111-rich matrix but not type I collagen matrix, indicating a LM-specific LaNt α31 effect(Troughton *et al.*, 2020b).

**Figure 1.**
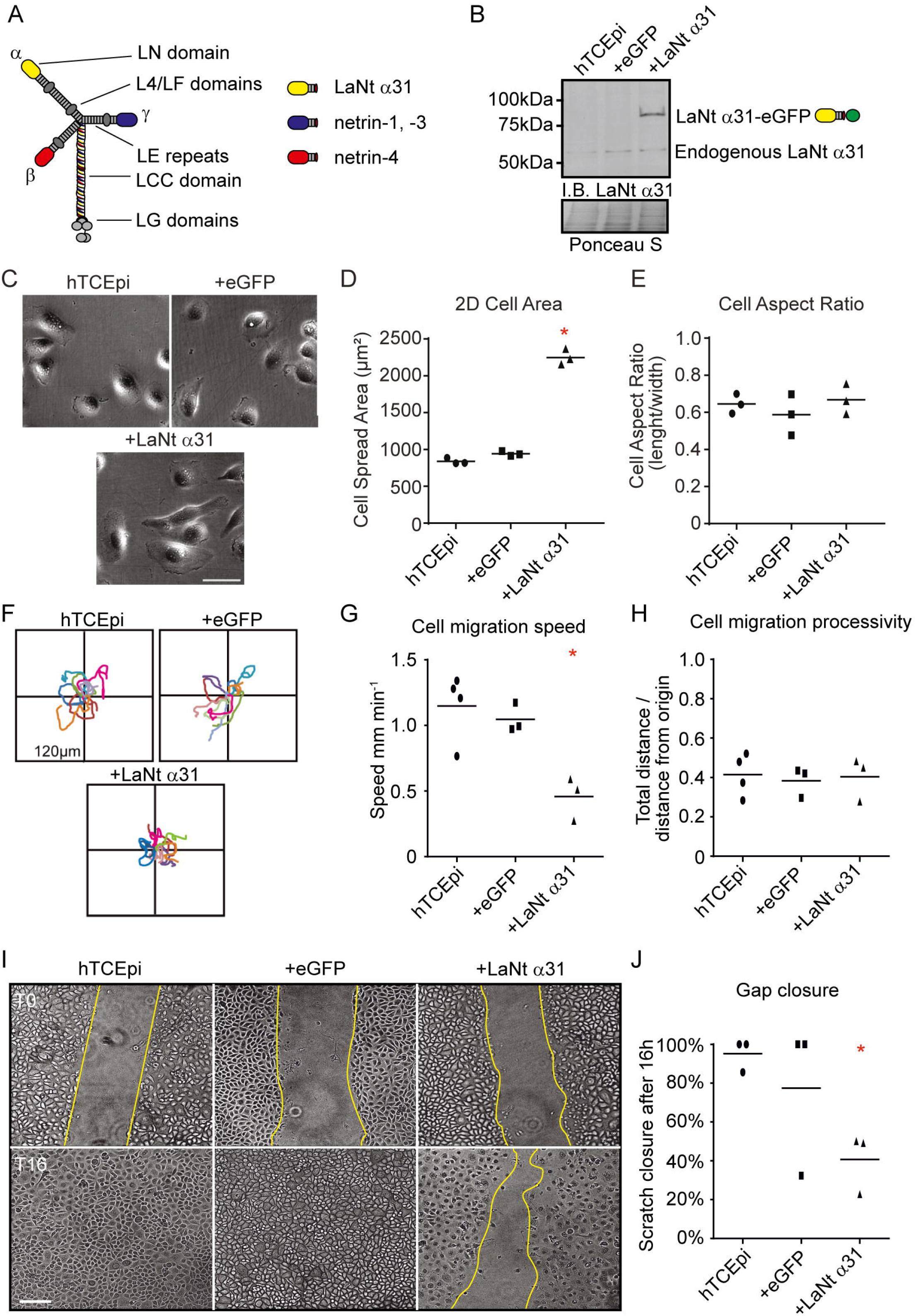
Corneal epithelial keratinocytes induced to express LaNt α 31 display decreased cell migration. (A) Diagrammatic representation of archetypal laminin, LaNt α31, and netrin protein structure. LN - laminin N-terminal domain, LE - laminin-type epidermal growth factor-like repeats. LCC - laminin coiled coil domain, LG - laminin globular domain. (B) Immunoblot of total cell lysates generated from hTCEpi cells transduced with either eGFP (+eGFP) or LaNt α31-eGFP (+LaNt α31) then probed with antibodies against LaNt α31. (C-H) Non-transduced (hTCEpi), +eGFP, or +LaNt α31 hTCEpi, were plated overnight onto tissue culture plastic at low density and either cell area and aspect area measured (C-E) or the migration paths of individual cells tracked over a 2 h period (F-H). (C) Representative phase contrast images, 2D cell area plotted in (D), and aspect ratio determined as length at longest point divided by width measured at 90° to that point (E). Each point represents one independent biological experiment with 40-60 cells per cell type, with line at mean. (F) Vector diagrams showing representative paths of 10 individual cells with each colour representing a single cell. (G) Migration speed measured as total distance migrated over time. (H) Migration processivity; measured as total distance migrated divided by maximum distance from the origin. In each graph lines are at mean with each point representing an individual experimental repeat consisting of the mean from 20-40 individual cells tracks. In (I) hTCEpi cells were plated overnight at confluence and a scratch wound introduced 16 h later. (I) Representative images from immediately after scratching (T0 upper panels) and after 16 h of recovery (T16 lower panels), yellow lines indicate wound margins. Scale bar 100μm. (J) Gap area closure measured 16 h after wounding plotted as percentage of the initial wound area with each point representing an independent experiment with either 2-3 technical repeats per experiment. * denotes p<0.05 compared with controls determined by one way ANOVA followed by Bonferroni post hoc test.

This simple hypothesis of LaNt α31 disrupting LM networks becomes more complex in epithelial tissues, including the ocular surface epithelium, where the most abundant LM is LM α3aβ3γ2 (LM332). LM332 differs from other LMs in that the α3a and γ2 subunits do not contain LN domains. As a consequence of this, LM332 is unable to polymerise independently *in vitro*(Schittny and Yurchenco, 1990; Cheng *et al.*, 1997). Despite this, LM332 is essential for the integrity of epithelial BMs(Pfendner and Lucky, 1993; Kivirikko *et al.*, 1995; Nakano *et al.*, 2002) and for epithelial wound repair(Goldfinger *et al.*, 1999) (Cameron *et al.*, 1988; Latvala *et al.*, 1995; Suzuki *et al.*, 2000). LM332 involvement in epithelial integrity is dependent on hemidesmosome (HD) cell-to-matrix adhesions, where the epithelial cells’ keratin intermediate filaments are anchored to LM332 *via* a protein complex containing the transmembrane proteins α6β4 integrin and type XVII collagen, the intracellular plakins plectin and bullous pemphigoid antigen 1e (BPAG1e) (Hopkinson *et al.*, 1998; Koster *et al.*, 2003; Hopkinson *et al.*, 2014) (Walko *et al.*, 2015). Formation of HDs proceeds in a step-wise manner with the recruitment of type XVII collagen and BPAG1e occurring only once nascent assemblies mature(Hopkinson *et al.*, 1998; Koster *et al.*, 2003), and maturation is associated with proteolytic processing of the LG 4-5 domains on LMα3 chain(Goldfinger *et al.*, 1999; Baudoin *et al.*, 2005). HD proteins are also involved in regulating cell migration(Frank and Carter, 2004; Pullar *et al.*, 2006; Sehgal *et al.*, 2006; Hamill *et al.*, 2009a; Hamill *et al.*, 2011); however, a second class of adhesive complex, focal adhesions (FAs), are considered the primary migratory force-generating machinery through providing linkage from LM332 to the actin cytoskeleton(Tsuruta *et al.*, 2011). Hundreds of different proteins are associated with FAs in different contexts; however, at the core of LM332 bound adhesions is the adaptor protein paxillin, which plays a scaffolding role through supporting recruitment of multiple other proteins(Ofuji, 2000; Turner, 2000). There is extensive cross-talk between HDs and FAs and recently it has been shown that this cross-talk is also relevant for force generation(Hopkinson *et al.*, 2014; Wang *et al.*, 2020).

Here we investigated the mechanism behind LaNt α31 influences upon corneal epithelial cell migration and adhesion, identifying that increasing expression of the LaNt α31 protein changes LM332 organisation and proteolytic processing, increased matrix metalloproteinase activity (MMP) and leads to assembly of mature HD-like complexes and altered FA distribution.

## Results

### High LaNt α31 expression curtails epithelial keratinocyte migration at times of new matrix synthesis

Previously, we reported that increased expression of LaNt α31 led to increased cell spreading and reduced migration of primary limbal-derived corneal epithelial cells (Barrera *et al.*, 2018). To investigate the mechanism for these effects we turned to the widely-used and well characterised limbal-derived corneal epithelial cell line hTCEpi (Yamamoto *et al.*, 2005; Dreier *et al.*, 2012; Yanez-Soto *et al.*, 2015; Hampel *et al.*, 2018; Kaplan *et al.*, 2019) (Supplemental Figure 1A) and used the same adenoviral construct to drive expression of LaNt α31 with a C-terminal eGFP tag (hereafter +LaNt α31, Figure 1B), comparing results against cells expressing eGFP alone (+eGFP). Consistent with previous findings in primary epithelial cells, +LaNt α31 hTCEpi mean 2D area was over twice as large as control cells when plated on uncoated tissue culture plastic (Figure 1, C and D, non-transduced hTCEpi 839 μm^2^ SD= ± 41 μm^2^, +eGFP 943 ± 33 μm^2^, +LaNt α31 2250 ± 110 μm^2^, p<0.05). No significant differences were observed in length/width aspect ratio (Figure 1E, non-transduced hTCEpi 0.65 ± 0.05, +eGFP 0.59 ± 0.11, +LaNt α31 0.67 ± 0.08, LM 0.65 ± 0.09). Also consistent with primary corneal epithelial cells, the +LaNt α31 hTCEpi cells displayed large reductions in total cell migration speed compared with controls (Figs. 1F, G, non-transduced hTCEpi 1.15 μm/min SD= ± 0.26, +eGFP 1.05 ± 0.11 μm/min, +LaNt α31-eGFP 0.46 ± 0.17 μm/min, p<0.05), with no significant differences in processivity, the ratio of total distance migrated to linear distance (Figure 1H, non-transduced hTCEpi 0.41 SD= ± 0.11, +eGFP 0.38 ± 0.08, +LaNt α31 0.40 ± 0.11). Cell sheet migration in gap closure assays also demonstrated a striking rate reduction in +LaNt α31 cell compared to control cells (Figs. 1I, 1J, closure after 16 h ± SD; non-transduced hTCEpi 95 ± 8 %, +eGFP 78 ± 39 %, +LaNt α31-eGFP 41 ± 16 %, p<0.05). Similar gap closure reduction effects were also observed in the HaCaT epidermal-derived keratinocyte and BHY oral epithelial cell line (closure after 16 h ± SD; non-transduced HaCaT 99 ± 2 %, +eGFP 97 ± 6 %, +LaNt α31 52 ± 12%, p<0.05; BHY 100 ± 0%, +eGFP 100 ± 0 %, +LaNt α31 88 ± 3 %, p<0.05. Supplemental Figure 1B, 2, A and B.).

### LaNt α31 interacts with LM332 and modifies its organisation

Having confirmed that hTCEpi cells represent an appropriate model, we next tested our hypothesis that LaNt α31 influences the LM ECM. First, the LM expression profile of hTCEpi, HaCaT and BHY cells was determined using RT-qPCR. For hTECpi, LAMA3A accounted for >95 % of the LM α chain encoding transcripts, LAMB3 >75 % of the β chains, and LAMC2 >85 % of the γ chains, indicating that LM3A32 is the predominant LM produced by these cells, with similar findings observed in HaCaT and BHY keratinocytes (Supplemental Figure 3). Immunoblotting of total protein extracts from non-transduced, +GFP, or +LaNt α31 hTCEpi revealed that induced LaNt α31 expression resulted in no major differences in absolute abundance of the expressed LMs, although an increased proteolytic processing of LMα3 was noted (Figure 2A).

**Figure 2.**
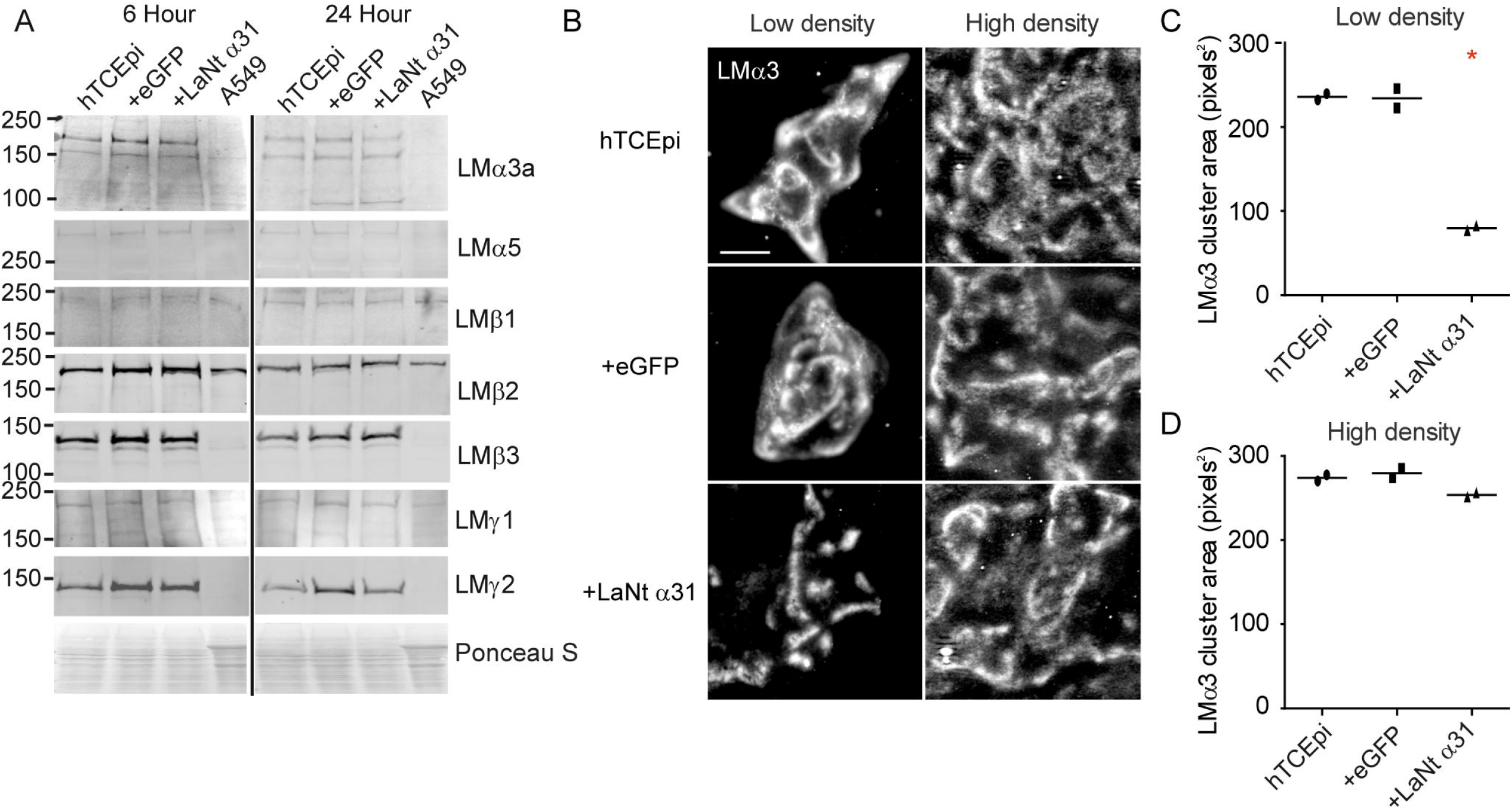
LaNt α31 expression induces changes to LM α3 organisation. (A) Immunoblots of total protein extracts from non-transduced, +eGFP, or +LaNt α31 hTCEpi, or A549 cells processed with antibodies against the indicated LM chain. Molecular weight markers are indicated to the left. Ponceau S staining was used to control for loading. (B) Non-transduced, +eGFP, or +LaNt α31 hTCEpi were seeded at low (left panels) or high (right panels) density on uncoated glass coverslips then processed for immunofluorescence microscopy with antibodies against LMα3 after 5 h (left panels) or 24 h (right panels). Scale bar 20 μm. (C) LMα3 cluster area was measured in non-transduced, +eGFP, or +LaNt α31 hTCEpi plated at low (C) or high (D) density. N=2 independent biological replicates; 25 measurements per cell type per assay. * In (C) denotes p<0.05 between +LaNt α31 and both controls, determined by one-way ANOVA followed by Tukey’s post-hoc test.

Next LMα3 organisation was investigated by indirect immunofluorescence microscopy with antibodies against LMα3. A striking change was observed in the LM organisation in the +LaNt α31 cells compared with controls. Specifically, whereas LMα3 was organised in broad arcs and contiguous lines in control cells, the LMα3 matrix deposited by +LaNt α31 cells was instead organised into tight, small clusters (Figure 2B, quantified in C; average LMα3 cluster area non-transduced hTCEpi 236 ± 5 pixel^2^, +eGFP 234 ± 16 pixel^2^, +LaNt α31 80 ± 4 pixel^2^, p<0.05). The same LMα3 clustering effect was also observed in primary corneal epithelial cells (Supplemental Figure 4, A and B). LMα3 signal co-distributed with LMβ3 (Supplemental Figure 4C), and the same clustering effect was observed using antibodies against LMβ3, LMγ2 and polyclonal antibodies against LM332 (Supplemental Figure 4D, E and F). However, no clear distribution pattern was observed for LMα5 in these cells irrespective of LaNt α31 expression, and together with the expression levels, this indicates the LaNt α31 effect was primarily upon LM332 (Supplemental Figure 4F). However, the phenotype was not apparent at higher cell densities when cells were allowed longer to depost LM, suggesting that LaNt α31 major influence is on the early stages of matrix assembly (Figure 2B, quantified in D; non-transduced hTCEpi 274 ± 5 pixel^2^, +eGFP 280 ± 8 pixel^2^, +LaNt α31 254 ± 3 pixel^2^).

We next asked where the LaNt α31-eGFP protein localised in the hTCEpi cells. Imaging live hTCEpi cells indicated a diffuse distribution of intracellular LaNt α31-eGFP throughout the cell; however, total internal reflection microscopy (TIRF) revealed organised clusters of LaNt α31-eGFP along the matrix apposed surface of the cells (Supplemental Figure 5A). The LaNt α31-eGFP signal remained present in ECM preparations after hyperosmotic removal of cellular material (Supplemental Figure 5B) and was similar to that reported when HaCaT ECM was processed using antibodies against LaNt α31 and LM332 (Hamill *et al.*, 2009c).

To investigate LaNt α31 effects on LM332 in live cells, an mCherry tagged LMβ3 adenovirus was used. Indirect immunofluorescence microscopy confirmed that the signal from the LMβ3 construct co-distributed with LMα3 and LMγ2 in hTCEpi indicating that it is incorporated into LM332 (Supplemental Figure 6). Imaging of live cells by TIRF microscopy revealed that although the LaNt α31-eGFP and LMβ3-mCherry signals were not identical, there were distinct regions where co-distribution was evident (Figure 3A). Moreover, when ECM extracts were prepared from fixed cells, the matrix-associated fluorescent LaNt α31-eGFP signal partially codistributed with LMβ3-mCherry (Figure 3B).

**Figure 3.**
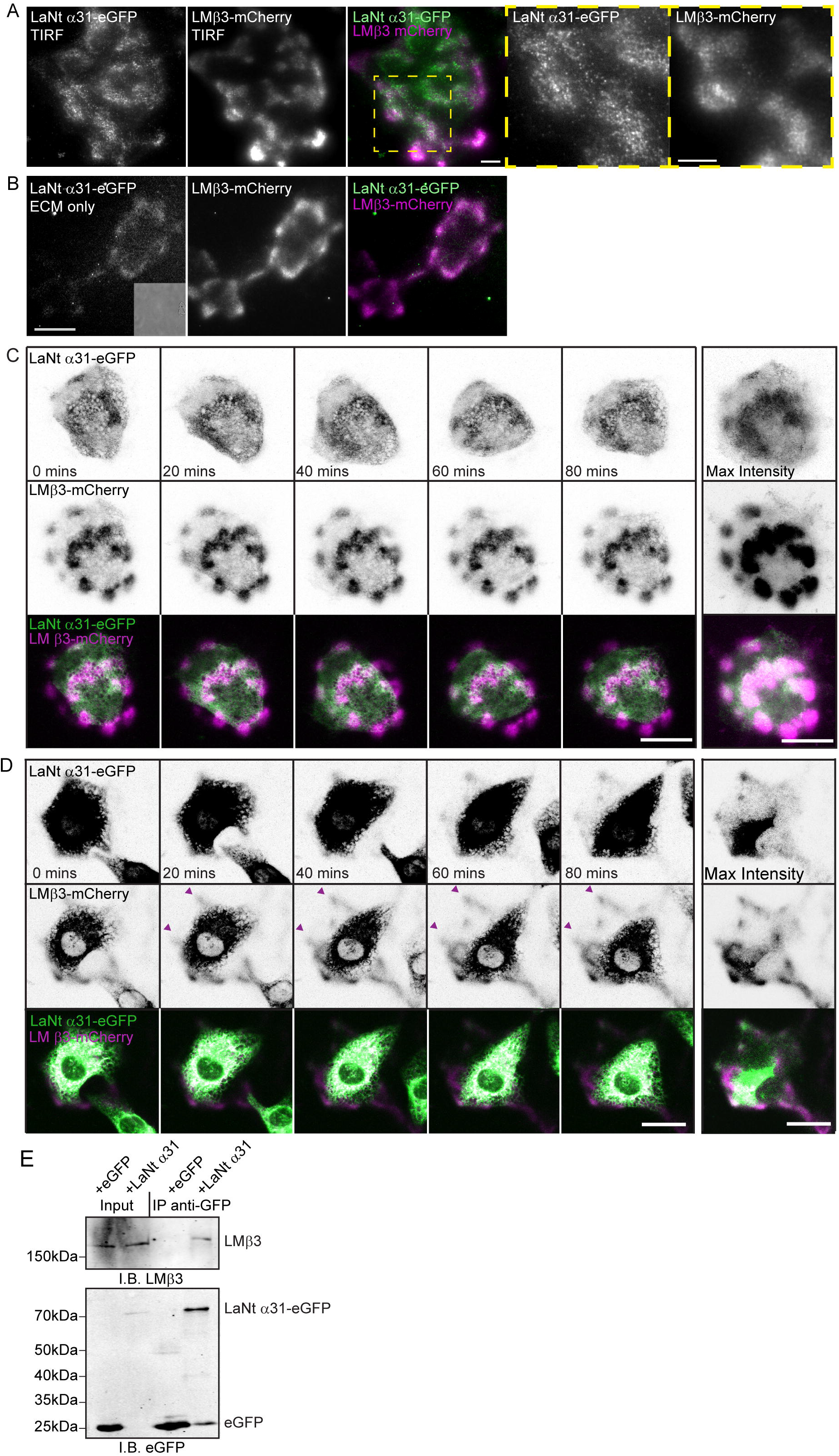
LaNt α31 co-distributes with LM β3. (A) TIRF images of hTCEpi cells doubly transduced with LaNt α31-eGFP (left panel, pseudocoloured green in merge) and LMβ3-mCherry (second panel, pseudocoloured magenta in merge) imaged 16 h after plating on glass-bottomed dishes. Yellow boxed region is shown at higher magnification in panels to the right. (B) ECM preparation from doubly transduced cells generated by ammonium hydroxide mediated removal from cells 16 h after plating, phase contrast image in inset shows absense of cellular material. Area imaged equates matrix deposited by approximately 1-3 cells (C and D) hTCEpi doubly transduced with LaNt α31-eGFP and LMβ3-mCherry were plated overnight (C) or for 2 h (D) on uncoated glass-bottomed dishes then imaged by confocal microscopy every 20 min over 16 h. (C and D) Representative images from 0 to 80 min of LaNt α31-eGFP (upper panels) and LMβ3-mCherry (middle panels), with signals inverted to aid visualisation. Bottom panel, merged images of LaNt α31-eGFP (pseudocoloured green) and LMβ3-mCherry (pseudocoloured magenta). Panels to far right are maximum intensity projection of the entire 16 h time course images. (E) Total cell lysates from hTCEpi transduced with eGFP (+eGFP) or LaNt α31-eGFP (+LaNt α31) were processed with anti-GFP antibody-conjugated beads. Input lysates equivalent to 8% of the total immunoprecipitation volume (left two lanes) and anti-GFP pull-down lanes (IP-anti-eGFP, right two lanes) were immunoblotted (IB) with antibodies against LMβ3 (upper panel) or GFP (lower panel). Scale bars in (A-D) 20 μm.

To investigate the protein dynamics of LaNt α31-eGFP relative to LMβ3-mCherry, cells were plated for 16 h on uncoated glass-bottomed dishes then imaged every 20 min over 12 h (Figure 3C, Supplemental Movie 1). Throughout this time course, the LaNt α31-eGFP rapidly formed and dissociated clusters at sites of LMβ3-mCherry deposits (Figure 3C). Moreover, over the first 3 h after plating cells onto uncoated dishes, i.e. during new matrix synthesis. The LMβ3-mCherry and LaNt α31-GFP proteins displayed similar patterns toward cell peripheries which could not be temporally resolved (Figure 3D). To determine whether the LaNt α31 - LMβ3 co-distribution represents formation of a complex containing the two proteins, we immunoprecipitated either eGFP or LaNt α31-eGFP then probed with antibodies against the endogenous LMβ3 (Figure 3E). LMβ3 precipitated along with LaNt α31-eGFP but not when eGFP alone was pulled-down (Figure 3E).

### Increasing LaNt α31 expression leads to mislocalised focal adhesions and early maturation of hemidesmosomes

Next, we used indirect immunofluorescence microscopy to analyse the effect of induced LaNt α31 expression on the two major cell-to-LM adhesive complexes: FA and HDs. This revealed that while paxillin formed discrete puncta in control hTCEpi cells, FAs in the +LaNt α31 cells were assembled into linear regions around cell margins (Figure 4A). As expected, the distribution of β4 integrin mirrored that observed for LM332, with swirls and arcs in +GFP control cells compared with tight clusters in the +LaNt α31 cells (Supplemental Figure 7).

**Figure 4.**
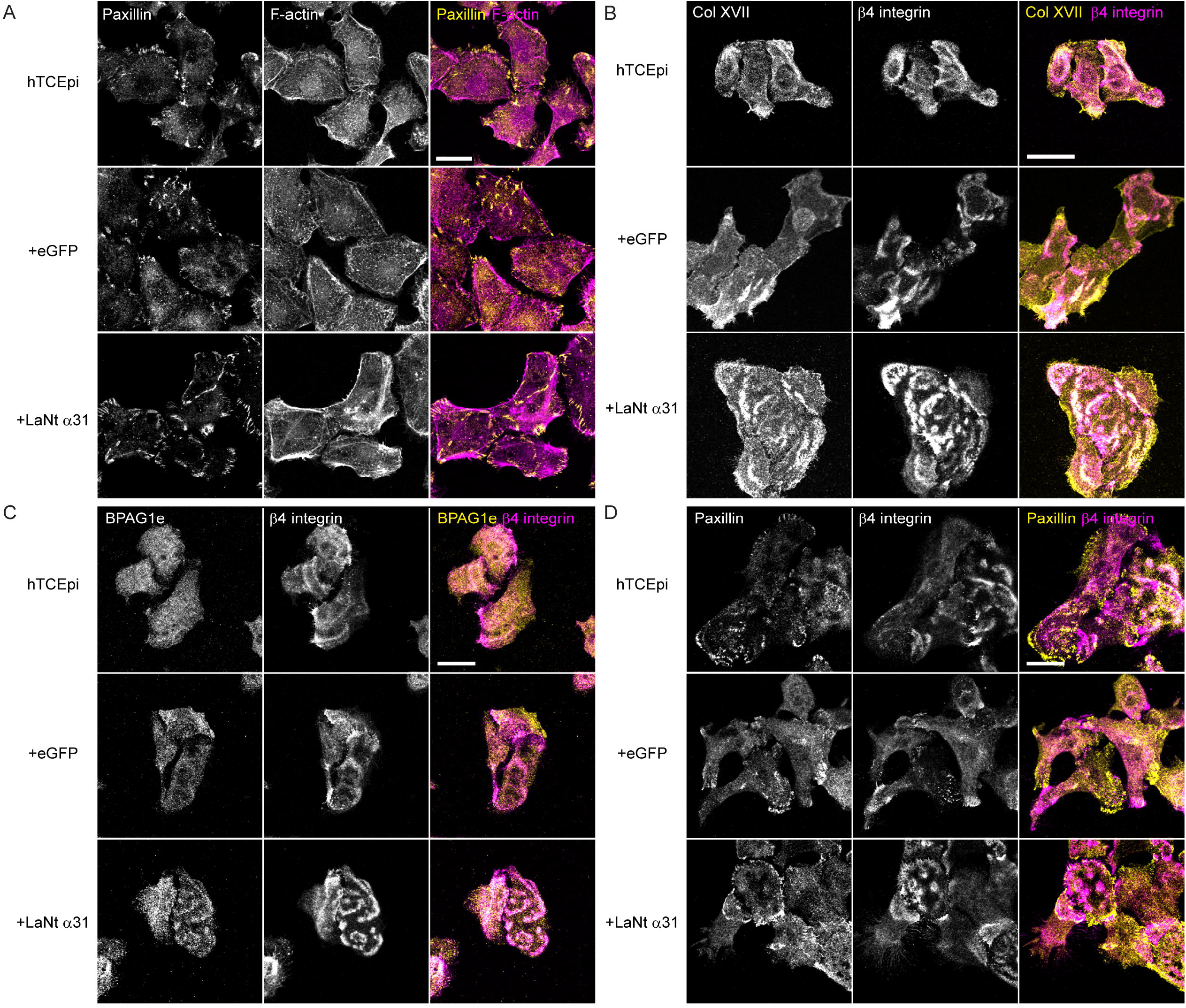
Focal adhesions are mislocalised and hemidesmosomes are more mature in LaNt α31 expressing corneal epithelial cells. Non-transduced, +eGFP, or +LaNt α31 hTCEpi were plated on glass coverslips then fixed and processed for indirect immunofluorescence with antibodies against: (A) paxillin (first column, yellow in merge) and with phalloidin to label filamentous actin (F-actin, second column, magenta in merge). (B) β4 integrin (first column, magenta in merge) and type XVII collagen (Col XVII, second column, yellow in merge). (C) bullous pemphigoid antigen 1e (BPAG1e, first column, yellow in merge) and β4 integrin (second column, magenta in merge). (D) paxillin (first column, yellow in merge) and β4 integrin (second column, magenta in merge). Scale bars 20 μm.

The later stages of HD maturation involve recruitment of type XVII collagen and BPAG1e to β4 integrin(Hopkinson *et al.*, 1998; Koster *et al.*, 2003). Immunofluorescence analysis of the distribution of type XVII collagen or BPAG1e in comparison to β4 integrin revealed increased colocalisation in +LaNt α31 cells compared with a general lack of co-distribution in non-transduced or +eGFP hTCEpi at this time point (Figure 4, B and C. Manders’ correlation coefficient type XVII collagen with β4 integrin: non-transduced hTCEpi 0.07 ± 0.12, +eGFP 0.22 ± 0.17, +LaNt α31 0.47 ± 0.19, p<0.05 1-way ANOVA with Bonferoni post-hoc test). Co-staining for paxillin and β4 integrin showed that the linear paxillin assemblies surrounded patches of β4 integrin (Figure 4D).

### Induced LaNt α31 expression increases matrix metalloproteinase activity

HD maturation is associated with proteolytic removal of the LG4/5 domains of LMα3, and analysis of total protein extracts had suggested that this was increased in +LaNt α31 cells (Figure 5A). To assess this further, ECM protein extracts were immunoblotted for LMα3 and the ratio of the processed ~160kD to the unprocessed ~190kDa band determined by densitometry. In the +LaNt α31 cells the processed proportion was almost two times higher compared with control-treated cells (Figure 5B, mean ± SD 160kDa to 190kDa ratios: non-transduced hTCEpi 0.4 ± 0.2, +eGFP 0.7 ± 0.2, +LaNt α31 1.3 ± 0.5, p=<0.05). Next, gelatin zymography was used to identify which MMPs are active in conditioned media from hTCEpi culture. This showed bands of gelatin degredation at molecular weights consistent with MMP2 and MMP9, consistent with reports in the literature (Daniels *et al.*, 2003; Pal-Ghosh *et al.*, 2011a) moreover, the zymography suggested that there could also be increased activity of these enzymes in +LaNt α31 cells (Figure 5C). This increased activity was then directly assesed using an MMP-specific fluorogenic substrate (Dnp-Pro-Leu-Gly-Leu-Trp-Ala-D-Arg-NH_2_ (Dnp=2,4-dinitrophenyl)) incubated with conditioned media from +LaNt α31 or control cells, measuring the fluorescence intensity over time (Figure 5D) These analyses revealed markedly increased activity in the +LaNt α31 cells: (mean+SD gradient of fluorescence emission curve: non-transduced hTCEpi 750 ± 70, +eGFP 870 ± 98, +LaNt α31 2980 ± 110, p=<0.05. Line of best r^2^ hTCEPi 0.84, +GFP 0.78, +LaNt 0.97). To determine if MMP activity was required for the LM clustering and HD maturation phenotype, +LaNt α31 hTCEpi cells were treated with either a hydroxamate-based broad spectrum MMP inhibitor (MMPi, CT-1746) or a mixture of inhibitors for serine, cysteine and aspartic proteases (non-MMPi, P1860, Figure 5E) and LMα3 organisation and HD maturation assessed by indirect immunofluorescence microscopy (Figure 5, F and G). The clustered LMα3 phenotype was evident in +LaNt α31 hTCEpi cells and in the +LaNt α31 cells treated with non-MMP inhibitors but was rescued in those cells treated with MMP inhibitor (Figure 5F). Similarly, the increased β4 integrin/ type XVII collagen, and β4 integrin/BPAG1e co-distribution, and the paxillin re-distribution observed in the +LaNt α31 cells were all rescued upon MMP inhibitor treatment but not with the non-MMP inhibitors (Figure 5G, Supplemental Figure 8, wA and B). Together these findings demonstrate that LaNt α31 requires MMP activity to exert its effects on LM organisation and HD maturation.

**Figure 5.**
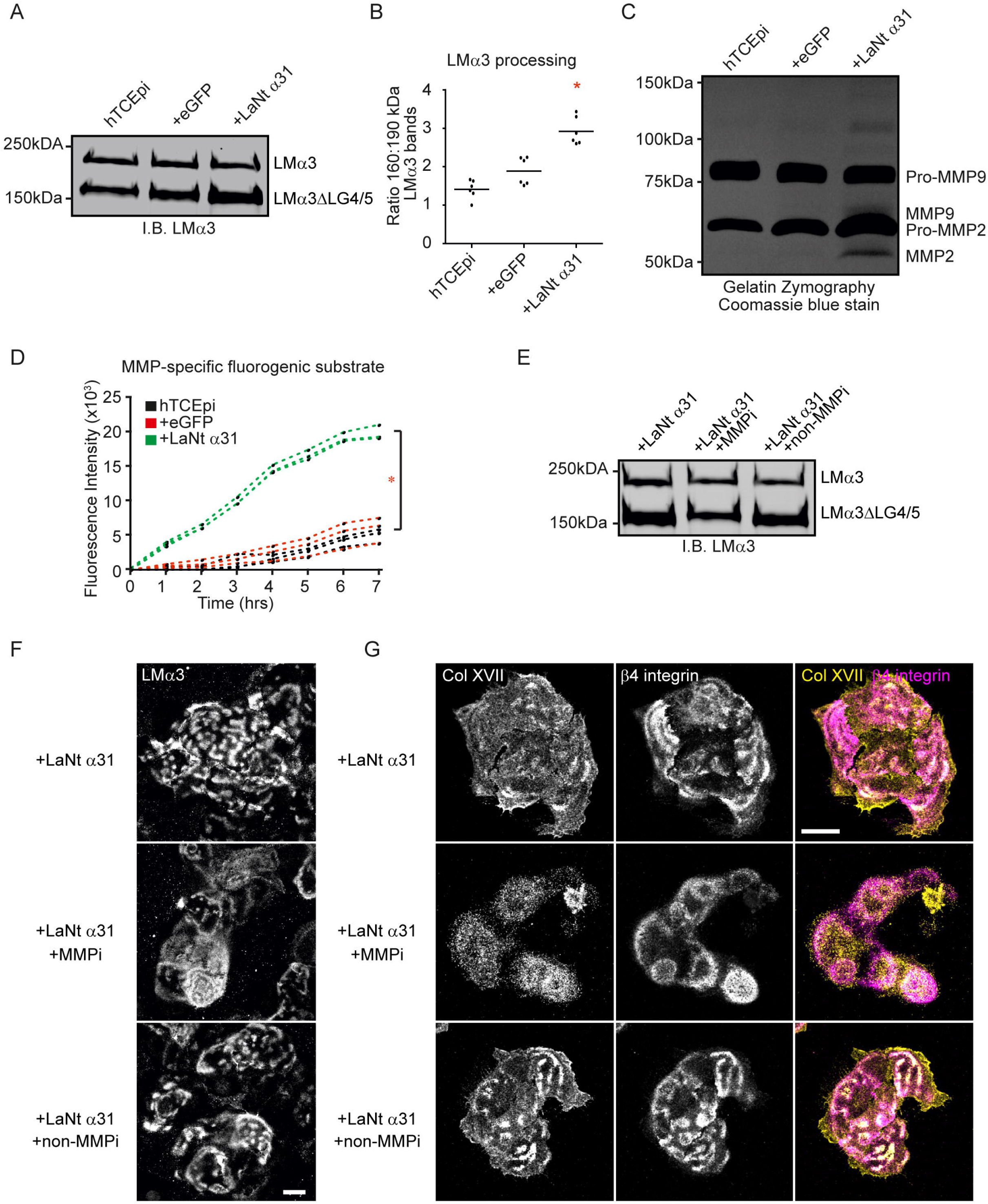
Induced expression of LaNt α31 increases MMP activity, which is required for LM α3 clustering and hemidesmosome maturation effects. (A and B) Non-transduced, +eGFP, or +LaNt α31 hTCEpi were plated overnight on tissue culture plastic and ECM extracts prepared by ammonium hydroxide-mediated hyperosmotic removal of cellular material. (A) Representative immunoblot with antibodies against LMα3. (B) Ratiometric quantification of 160 kDa band to 190 kDa band. Each point represents an independent experiment. * denotes p<0.05 compared with both controls as determined by one-way ANOVA with Bonferroni post-hoc test. (C) Gelatin zymography of conditioned media extracts generated from non-transduced, +eGFP, or +LaNt α31 hTCEpi. Indicated to the right of the gel image are the predicted sizes of pro and active forms of MMP2 and 9. (D) Conditioned media extracts were normalised for cell number then incubated with the BML-P131-0001 peptide, which emits fluorescence only once cleaved by MMPs. The graph displays the fluorescence intensity measured over time with each line representing an independent experiment. (E and F) +LaNt α31 hTCEpi cells were plated overnight with broad spectrum MMP inhibitor (MMPi) or serine protease inhibitor (non-MMPi) and extracellular matrixes prepared for western blotting (E) or coverslips processed for indirect immunofluorescence microscopy with antibodies LMα3 (F) or type XVII collagen and β4 integrin (G). Scale bar 20 μm. * In (D) denotes p<0.05 between +LaNt α31 and both controls, determined by ACOVA..

### Provision of preformed matrix rescues the LaNt α31 induced cell spreading and motility phenotypes

All the results to this point supported the hypothesis that LaNt α31 influences LM matrixes. However, we could not rule out the effects being driven at the cellular level. Therefore, we asked whether provision of a cell-derived ECM from hTCEPi cells could rescue the spreading and migration phenotypes (Figure 6A). Cell-derived ECMs from non-transduced hTCEpi (hTCEpi ECM) and +LaNt α31-eGFP hTCEpi (+LaNt α31 ECM) were generated from cells plated at ~90% confluence for 24 h. Onto these prepared matrices fresh hTCEpi or +LaNt α31 cells were plated and 2 h later the cell spreading (Figure 6, B-D) and migration (Figure 6, E-G) were analysed. These studies revealed that plating +LaNt α31 cells upon the matrix from hTCEpi cells rescued both the spreading and cell migration phenotypes with no statistically significant differences between hTCEpi and +LaNt α31 cells (cell spreading: hTCEpi on hTCEpi ECM 1180 ± 280 μm^2^ compared with +LaNt α31 on hTCEpi ECM 1300 ± 280 μm^2^, cell motility, hTCEpi on hTCEpi ECM 1.16 ± 0.08 μm/min, +LaNt α31-GFP on hTCEpi ECM 1.20 ± 0.05 μm/min,).

**Figure 6.**
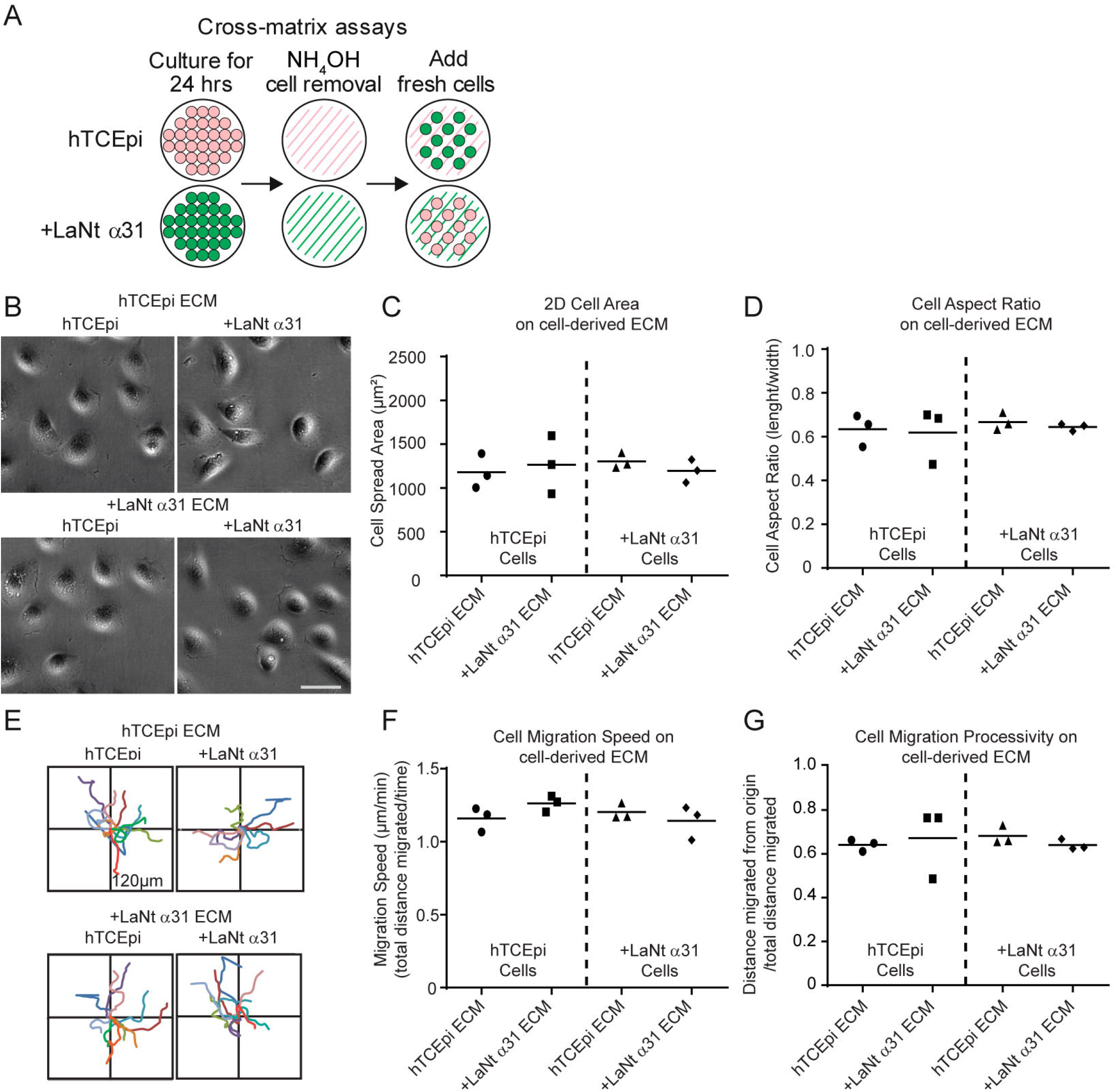
LaNt α31 overexpression adhesion and migration defects are rescued through provision of a cell-derived matrix. (A) Diagramatic representation of cross-matrix assays. Non-transduced or +LaNt α31 hTCEpi were plated overnight at high density and cell-derived matrices prepared by ammonium hydroxide removal of cellular material. Onto prepared matrixes, fresh non-transduced or +LaNt α31 hTCEpi were seeded at low density, allowed to adhere for 2 h then imaged. (B) Representative phase contrast images. (C) 2D cell spread area and (D) aspect ratio determined by measuring 40-60 cells per cell type per experiment with each point on the graph representing one independent biological experiment with line at mean. (E) Representative vector diagrams of migration paths of 10 randomly selected cells tracked over a 2 h period. (F) Migration speed and (G) processivity with each point representing an individual experimental repeat consisting of the mean from 20-40 individual cells tracks.

The matrix deposited by high-density +LaNt α31 cells was also indistinguishable from hTCEpi ECM in terms of supporting cell movement of the non-transduced hTCEpi cells and the +LaNt α31 cells (spreading; hTCEpi on +LaNt α31 ECM 1270 ± 230 μm^2^, +LaNt α31 on +LaNt α31 ECM 1190 ± 300 μm^2^, migration; hTCEpi on +LaNt α31 ECM 1.26 ± 0.05 μm/min, +LaNt α31 on +LaNt α31 ECM 1.14 ± 0.12 μm/min. Figure 6, B-G). These data are in agreement with the analysis of LMα3 organisation from high-density cultures being indistinguishable for control cultures (Figure 2D) and indicate that the +LaNt α31 effects are restricted to times when new matrix deposition is required.

## Discussion

The findings presented here have identified that LaNt α31 forms a complex and is depositied together with LM332, and during times of new ECM deposition, alters LM332 organisation and proteolytic processing. These changes were associated with increased MMP activity and clustering of deposited LM, changes to FA distribution and maturation HD-like complexes. Together these findings implicate the LaNt protein family as new regulators of LM-dependent cellular processes. Together, with previous findings indicating changes to LaNt α31 expression during corneal wound repair(Barrera *et al.*, 2018), these results suggest that controlling LaNt α31 levels represents a new mechanism to regulate the switch from a migratory phenotype to stably attached epithelial sheets once reepithelialisation has occurred. This susggests that LaNt α31, and by extension, the other LaNt family members, may help to define the “Goldilocks” level of cell-matrix adhesion required for specific epithelial cell behaviours.

The chicken vs egg question of whether the MMP and FA/HD changes are a result of the changed LM organisation or *vice versa* is particularly challenging to resolve at this point. This is not least because extensive feedback occurs between these processes. The tighter and more restricted LM332 deposition pattern supporting the maturation of HD complexes suggests the possibility that the additional α LN domain from LaNt α31 provides partial stabilisation of weak LM-LM interactions that could not usually occur. However, according to the robustly established models of LN-LN interactions(Yurchenco and Cheng, 1993), in the LM3A32-rich matrix deposited by hTCEpi cells, the absence of a γ LN domain in LM332 means that any LaNt α31 interaction with LMβ3 would, at best, be unstable (Hussain *et al.*, 2011; Carafoli *et al.*, 2012)(Schittny and Yurchenco, 1990; Yurchenco and Cheng, 1993; Cheng *et al.*, 1997; Hussain *et al.*, 2011; Carafoli *et al.*, 2012). Epithelial cells are known to also produce small amounts of LM511, 521 and 111 (Ljubimov *et al.*, 1995; Kabosova *et al.*, 2007; Schlotzer-Schrehardt *et al.*, 2007), which is consistent with our RT-qPCR and immunoblotting data. Each of these additional LMs provide α, β and γ LN domains and it is possible that the observed LM332 clusters contain some of these additional LMs, although we have not been able to detect them by indirect immunofluorescence microscopy. Overall, although the data support some form of LaNt α31–LM332 interactions, we think it unlikely that it is a direct interaction driving the observed effects.

All the LaNt α31 induced effects were observed at times where new matrix deposition was required. LM assembly and organisation requires receptor binding, driven predominantly by high-affinity binding to sites within the LM LG domains. In turn, the engagement of LM receptors drives further LM recruitment. One model our data could support is that LaNt α31 to receptor binding helps to initiates this cascade (Figure 7). In support of this model, cell-surface receptor-binding capabilities have been described for some LN domains(Garbe *et al.*, 2002; Odenthal *et al.*, 2004), although these affinities are lower than the LM-integrin binding interactions(Nishiuchi *et al.*, 2006; Taniguchi *et al.*, 2009; Taniguchi *et al.*, 2020). In the case of LaNt α31 and LM332, the spatial colocalisation and immunoprecipitation data indicate formation of a complex containing both proteins, which suggests that any receptor-mediated effects may require interaction. The concept of a LaNt α31/ LM332 complex mediating interactions is consistent with studies of netrin-4 which has been shown can regulate α6β1 integrin activity in a complex with LMγ1 (Staquicini *et al.*, 2009). Additionally, knockout α3 or β4 integrin both lead to changes to LM332 organisation, clearly indicating a direct role for FAs and HD in defining LM332 patterning (deHart *et al.*, 2003; Sehgal *et al.*, 2006). This model also supports the MMP activity increase: although we observed a large overall increase in MMP activity, this could be achieved through a modest initial effect as feedback from MMP-induced release of LMα3 LG4/5 has been shown to drive further MMP activation (Momota *et al.*, 2005; Michopoulou *et al.*, 2020).

**Figure 7.**
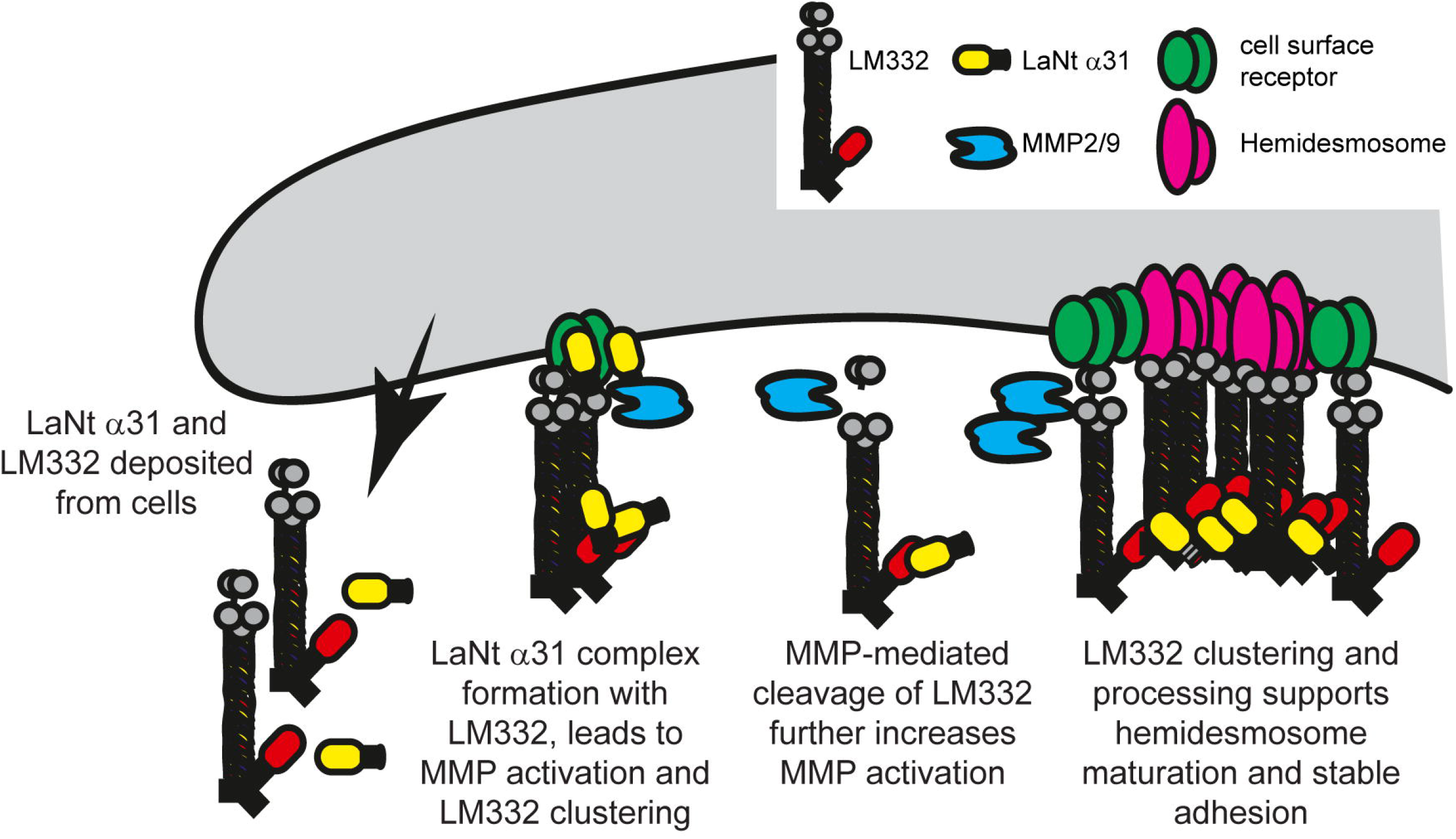
Model for LaNt α31 mediated effects.

Localised and temporally regulated MMP activity is required for effective wound repair but excessive or uncontrolled activity is deleterious for tissue homeostasis, including leading to complications following wound repair such as recurrent corneal erosions (Matsubara *et al.*, 1991a; Pal-Ghosh *et al.*, 2011a; Pal-Ghosh *et al.*, 2011b Matsubara, 1991 #101). Confirmation of our model would indicate that relatively modest changes in LaNt α31 expression could initiate larger changes to BM structure and function, with implications for many aspects of tissue function, while also raising the potential for therapeutic intervention.

The findings reported here demonstrate that LaNt α31 influences LM organisation, MMP-mediated LM processing, and cell-to-matrix adhesion. As the LaNt proteins are produced by alternative splicing from LM genes, these new data indicate that control of the LaNt to LM ratio adds a hitherto unknown layer of self-regulation of LM function. Combined with LaNt α31 protein being enriched in numerous adult tissues (Troughton *et al.*, 2020a), the findings detailed here suggest that this relatively unstudied protein and this new LM-autoregulatory mechanism could have widespread implications particularly for wound repair and other situations where BM remodelling is required, such as tissue morphogenesis and development.

## Methods

### Antibodies and other reagents

Rabbit monoclonal antibodies raised against paxillin (clone Y113), were used at 10 μg/ml for immunofluorescence microscopy (IF), mouse monoclonal antibodies raised against β4 integrin (clone M126, ab29042) were used at 4 μg/ml, and rabbit monoclonal antibodies raised against type XVII collagen (clone EPR18614, ab184996) were used at 10 μg mL^−1^ for IF and 0.25 μg/ml for immunoblotting (IB), all from Abcam (Abcam, Cambridge, UK). Mouse monoclonal antibodies for IF against LMα3 (clone RG13) (Gonzales *et al.*, 1999) used at 10 μg/ml, rabbit polyclonal antibodies raised against LM332 (J18) (Hopkinson *et al.*, 1992) and cytokeratin 12 {Kurpakus, 1990 #12636} were used at 1:50, and human monoclonal antibodies to BPAG1e (5E) (Hashimoto *et al.*, 1993) used at 1:2, were generous gifts from Jonathan Jones, Washington State University, WA. Mouse monoclonal antibodies raised against human LaNt α31 (clone 3E11) used at 1.9 μg/ml for IB were described previously (Barrera *et al.*, 2018; Troughton *et al.*, 2020a). Rat monoclonal antibodies raised against LMγ1 (clone A5, ab80580) used at 1 μg/ml, and mouse monoclonal antibodies raised against LMβ2 (clone CL2979) used at 1 μg/ml were from Abcam. Mouse monoclonal antibodies raised against LMα3 (clone CL3112, HPA001895) used at 0.5 μg/ml, vimentin (clone V9, V6389) used used at 0.01 μg/ml, LMα5 (clone 2F7, WH0003911M1) used at 1 μg/ml, and against GFP (a mixture of clones 7.1 and 13.1, 11814460001) used at 0.08 μg/ml, were from Sigma-Aldrich (Sigma-Aldrich, St. Louis, Missouri, USA). Rabbit polyclonal antibodies raised against LMβ1 (PA5-27271) were used at 1 μg/ml, against LMβ3 (PA5-29161) at 0.46 μg/ml, against LMγ2 (PA5-51110) at 0.66 μg/ml, Alexa Fluor™ 647 Phalloidin (A22287) used at 5 units/ml, were from ThermoFisher (ThermoFisher Scientific, Waltham, Massachusetts, USA). Alexa Fluor 594 nm, 647 nm or Cy5 conjugated goat anti-mouse and goat anti-rabbit secondary antibodies were obtained from Jackson Immunoresearch (Stratech, Ely, UK). Goat anti-mouse IRDye 800CW and goat anti-rabbit IRDye 680CW were obtained from LI-COR BioSciences (LI-COR BioSciences, Lincoln, Nebraska, USA).

For MMP inhibition, hydroxamate-based MMP inhibitor CT-1746 (N1-[2-(S)-(3,3-dimethylbutanamidyl)]-N4-hydroxy-2-(R)-[3-(4-chlorophenyl)-propyl]-succinamide) was diluted to 100 μM in culture media (UCB Celltech, Slough, U.K). For inhibition of serine, cysteine and aspartic proteases, P1860 inhibitor cocktail consisting of aprotinin, bestatin, E-64, leupeptin, pepstatin A was diluted to 1:1000 in culture media (Sigma-Aldrich).

### Cell Culture

Telomerase-immortalised human corneal epithelial cells, hTCEpi cells(Robertson *et al.*, 2005), were cultured at 37 °C with 5 % CO₂ in keratinocyte-serum-free medium (KSFM) supplemented with bovine pituitary extract (BPE, 0.05 mg/ml), human recombinant Epidermal Growth Factor (5 ng/ml, ThermoFisher Scientific) and 0.15 mM CaCl₂ (Sigma-Aldrich). HaCaT spontaneously transformed human epidermal keratinocytes(Boukamp *et al.*, 1988), BHY non-metastatic oral squamous carcinoma cells(Kawamata *et al.*, 1997) and A549 lung adenocarcinoma cells(Giard *et al.*, 1973) were cultured using Dulbecco’s Modified Eagle Medium (DMEM, Sigma-Aldrich) supplemented with 10 % fetal calf serum (LabTech, East Sussex, UK) and 4 mM L-glutamine (Sigma-Aldrich).

### Adenovirus production and cell transduction

Adenoviral constructs inducing expression of CMV-drive full-length human LaNt α31 with C-terminal eGFP tag were described previously(Barrera *et al.*, 2018). Adenoviruses inducing CMV driven expression of LMβ3-mCherry(Hopkinson *et al.*, 2008) and eGFP were kind gifts from Jonathan Jones, Washington State University, WA.

hTCEpi, HaCaT, or BHY cells were cultured in T75 flasks (Greiner-Bio One, Stonehouse, Gloucestershire, UK) until 70 % confluent or were seeded at 1 x 10^6^ cells/plate in 100 mm plates for 24h. Cells were then transduced with eGFP, LaNt α31-eGFP or LMβ3-mCherry adenoviruses. Transduction efficiency was confirmed by fluorescence microscopy or flow cytometiry as being 60-90% at point of use. Experiments were conducted 24-48 h following transduction.

### SDS PAGE, immunoblotting and immunoprecipitation

For total protein extracts, 1 x 10^6^ cells were seeded in 100 mm dishes (Greiner-BioOne) then lysed after 6 h or 24 h in urea/ sodium dodecyl sulfate (SDS) buffer (10 mM Tris-HCl pH 6.8, 6.7 M Urea, 1 % SDS, 10 % glycerol and bromophenol blue) containing 50 μM phenylmethylsulfonyl fluoride (PMSF) and 50 μM N-ethylmaleimide (all Sigma-Aldrich). For LMα3 ECM processing blots, cells were seeded at 2.5 x 10^5^ cells/well in 6-well plates. After 16 h, cellular material was removed through exposure to 0.18 % NH_4_OH for 5 min followed by extensive PBS washes (Langhofer *et al.*, 1993), and ECM solubilised in urea/ SDS buffer. Before loading, lysates were sonicated, and 10 % β-mercaptoethanol (final volume) added.

For immunoprecipitation 48 h after transduction cells were extracted in 0.1 % SDS, 0.5% sodium deoxycholate, 1 % Nonidet P-40, 150 mM NaCl, 1 mM CaCl_2_ in 50 mM Tris-HCl, pH 7.5 with protease and phosphatase inhibitors (all Sigma-Aldrich). Cell extracts were clarified by centrifugation, and a 50 % slurry of sepharose beads covalently conjugated with rabbit anti-GFP polyclonal antibodies (Abcam) added to the supernatant and incubated overnight at 4 °C. Beads were washed in lysis buffer, collected by centrifugation and boiled in SDS-PAGE sample buffer (50 mM Tris-HCl pH 6.8, 10 % glycerol, 1 % SDS, 10 % BME).

Proteins were separated by SDS-polyacrylamide gel electrophoresis (SDS-PAGE) using 7.5 % or 10 % polyacrylamide gels (1.5 M Tris, 0.4 % w/v SDS, 7.5 % 10 % acrylamide/ bis-acrylamide; electrophoresis buffer: 25 mM Tris, 250 mM glycine, 0.1 % w/v SDS, pH 8.5. All Sigma-Aldrich) and transferred to a nitrocellulose membrane using a Biorad TurboBlot system (BioRad, Hercules California, USA) or a BioRad wet transfer system (12.5 mM Tris, 125 mM glycine, 0.05% SDS, 20 % ethanol), and blocked for 1 hr at room temperature in Odyssey^®^TBS-Blocking Buffer (LI-COR BioSciences) or in 5 % (w/ v) Marvel skimmed milk in TBS (Tesco, Hertfordshire, UK). The membranes were probed with primary antibodies diluted in blocking buffer overnight at 4 °C, washed 3 x 5 min in TBS with 0.1 % Tween (TBS-T) then probed for 1 hr at room temperature with IRDye^®^ 800CW or IRDye^®^ 680CW conjugated secondary antibodies (LI-COR) diluted in blocking buffer (0.05 μg/ml). Membranes were then washed for 3 x 5 min in TBS-T and imaged using an Odyssey^®^ CLx 9120 Infrared Imaging System (LI-COR).

### Gap closure, cell migration assays, and cell morphology analyses

For gap closure assays, cells were seeded into ibidi® 2-well culture inserts (Ibidi, Martinsried, Germany); at 7.0 x 10^4^ cells/well. Culture inserts were carefully removed after 6 h, cell debris washed away, and the gap margin imaged using brightfield optics on a Nikon TiE epifluorescence microscope with a 10X objective at 0 and 16 h (Nikon, Tokyo, Japan). Gap closure was measured as a percentage relative to starting area using the freehand tool in Image J (NIH, Bethesda, Massachusetts, USA).

For cell morphology analyses and low-density migration assays, cells were seeded at 2.5 x 10^4^ cells/well onto uncoated 24-well plates (Greiner-Bio One). Where cell-derived preformed matrixes were required, non-transduced hTCEpi or +LaNt α31 hTCEpi were plated at 1.8 x 10^5^ cells/well of a 24-well plate and cultured overnight, after which cellular material was removed with 0.18 % NH_4_OH for 5 min followed by extensive PBS washes. Fresh untreated or +LaNt α31 hTCEpi were then seeded on top of the prepared matrixes and incubated for 2 h.

For morphology analyses, phase-contrast images were acquired on a Nikon TiE microscope and analysed using ImageJ software. Cell perimeters were manually traced to define cell area; cell length was measured as the longest linear axis, with width measured at widest point at right angles to the length measurement. The aspect ratio was defined as the ratio of width to length. For low-density migration assays, cells were imaged every 2 min over a 2 h period, using a 20X objective on a Nikon TiE fluorescent microscope, then individual cells tracked using the MTrackJ plugin on ImageJ. Speed (total distance travelled/time) and processivity (total distance/ distance from the origin) were calculated for each cell.

### Immunofluorescence microscopy

7.5 x 10^4^ (low density) or 1.5 x 10^5^ (high density) cells were seeded onto no. #1 round 16 mm glass coverslips (Pyramid innovations Ltd, Polegate, UK) in 12-well plates (Greiner-BioOne) and cultured overnight for approximately 16 h before fixing and extracting in ice-cold methanol for 15 min, or in 3.7 % formaldehyde (Sigma-Aldrich) for 10 min followed by 10 min in 0.2 % Triton X-100 in PBS to permeabilise (Sigma-Aldrich). For ECM-only analysis, cellular material was removed with 0.18 % NH_4_OH for 5 min followed by extensive PBS washes prior to fixation. Primary antibodies were diluted in PBS with 10% normal goat serum (Jackson Labs) and incubated at 4 °C overnight; coverslips were then washed 3x for 5 min with PBS containing 0.05 % tween-20 (PBS-T, Sigma-Aldrich) before probing for 1 h at room temperature with Alexa Fluor 594nm or 647nm/ Cy5 conjugated secondary antibodies diluted in PBS. Coverslips were washed 3x for 5 min with PBS-T, counterstained with DAPI (ThermoFisher Scientific) for 10 min, rinsed thoroughly with deionised H_2_0 and mounted with Vectashield (Vector Laboratories, Burlingame, California, USA). Images were obtained using a Zeiss LSM800 confocal microscope (Zeiss, Cambridge, UK).

LMα3 staining area was measured for 50 regions of staining per cell treatment using the freehand selection tool on ImageJ. For type XVII collagen colocalisation with integrin β4, Manders’ correlation coefficients were calculated using the Coloc 2 plugin on ImageJ for all pixels above channel thresholds.

### Live protein imaging

For TIRF microscopy and new matrix deposition assays, 1 x 10^5^ hTCEpi cells, doubly transduced with LaNt α31 GFP and LMβ3 mCherry adenoviruses were seeded onto uncoated 35 mm glass-bottomed dishes (MatTek Corporation, Ashland, Massachusetts, USA) then imaged after either 2 h or after 16 h using a Zeiss 510 Multiphoton 2 confocal microscope or Zeiss LSM880 confocal laser scanning microscope with TIRF capability. In live experiments, images were acquired using a 63X objective every 20 min over 16 h. Images were processed using ImageJ.

### MMP activity assays

hTCEpi cells were seeded at 8 x 10^5^ cells per dish in 60 mm dishes (Greiner-BioOne) and cultured in 2 mL BPE-free KSFM media for 48 h. Conditioned media was collected and concentrated to 1 mL using Amicon Ultra-0.5 centrifuge filters with 10 kDa pore size (Sigma-Aldrich). Cell counts were achieved by fixing the cells in 3.7 % formaldehyde for 10 min, staining with 0.25 μg/ml DAPI for 10 min (ThermoFisher Scientific), imaging at the centre of the dish using 4x objective on a Nikon TiE microscope, then counting using the analyse particles tool on image J. The final volumes of harvested media normalised for cell number, and supplemented with 0.1 % bovine serum albumin (BSA, Sigma-Aldrich) to stabilise MMPs.

For gelatin zymography assays, concentrated normalised conditioned media was mixed with SDS-PAGE sample buffer without BME on a 1 mg/ml gelatin-containing 10 % polyacrylamide gel (gelatine, Sigma-Aldrich). The gels were washed for 2 h in renaturing buffer (50 mM Tris-HCL containing 2.5 % Triton-x 100), and incubated at 37 °C in zymography buffer (50 mM Tris-HCL containing 5 mM CaCl_2_) for 24 h. The gels were then stained overnight at 4 °C, destained for 2 h at room temp (Coomassie Brilliant Blue R-250 staining kit, BIO-RAD), and imaged on a ChemiDoc MP (BIO-RAD).

For fluorogenic substrate assays, conditioned media was diluted 1:1 zymography buffer containing 10 μM peptide (BML-P131-0001, Dnp-Pro-β-cyclohexyl-Ala-Gly-Cys(Me)-His-Ala-Lys(Nma)-NH_2,,_ Enzo Lifesciences, Exeter, UK) and 10 mM CaCl_2_ (final conc. 5 μM/ 5 mM, respectively) and 100 μl added in triplicate to a black 96-well plates (Corning, Somerville Massachusetts, USA). Plates were read every hour for 7 h at an excitation/ emission spectra of 280 nm/ 350 nm on a SPECTROstar-nano plate reader (BMG LABTECH Ltd, Bucks, UK).

### Statistical analyses

Quantification of experiments was derived from a minimum of 3 independent experiment repeats. One-way ANOVA and Tukey’s HSD or Bonferroni post hoc analysis or linear regression and ANCOVA were used to analyse and interpret the data using GraphPad Prism 6 (GraphPad Software, San Diego, California, USA).

## Abbreviations

LM: laminin
LaNt: Laminin N-terminus proteins
LN: laminin N-terminal domain
LE: laminin-type epidermal growth factor-like
LCC: laminin coiled-coil domain
MMP: matrix metalloproteinase
HD: hemidesmosome
FA: focal adhesion
ECM: extracellular matrix
TIRF: total internal reflection microscopy
IF: immunofluorescence microscopy
IB: immunoblotting
DMEM: Dulbecco’s modified Eagle’s medium
KSFM: keratinocyte serum free media
BPE: bovine pituitary extract

## Acknowledgements

The authors would like to acknowledge Jonathan Jones and Susan Hopkinson (Washington State University) and Robert Lavker (Northwestern University) for generous gifts of reagents and cells. Technical help was greatly appreciated from David Mason, Marco Marcello, Violaine See and Joanna Majchrowska at the Centre for Cell Imaging at the University of Liverpool.

This work was supported by funding from the Biotechnology and Biological Sciences Research Council (BB/L020513/1), Fight For Sight (New lecturers’ award), British Skin Foundation (PhD studentship), Foundation for the Prevention of Blindness, and Versus Arthritis Career Development Fellowship (21447).

## References

Aumailley, M. (2013). The laminin family. Cell adhesion & migration 7, 48–55.

Barrera, V., Troughton, L.D., Iorio, V., Liu, S., Oyewole, O., Sheridan, C.M., and Hamill, K.J. (2018). Differential Distribution of Laminin N-Terminus alpha31 Across the Ocular Surface: Implications for Corneal Wound Repair. Investigative ophthalmology & visual science 59, 4082–4093.

Baudoin, C., Fantin, L., and Meneguzzi, G. (2005). Proteolytic processing of the laminin alpha3 G domain mediates assembly of hemidesmosomes but has no role on keratinocyte migration. The Journal of investigative dermatology 125, 883–888.

Boukamp, P., Petrussevska, R.T., Breitkreutz, D., Hornung, J., Markham, A., and Fusenig, N.E. (1988). Normal keratinization in a spontaneously immortalized aneuploid human keratinocyte cell line. The Journal of cell biology 106, 761–771.

Cameron, J.D., Hagen, S.T., Waterfield, R.R., and Furcht, L.T. (1988). Effects of matrix proteins on rabbit corneal epithelial cell adhesion and migration. Current eye research 7, 293–301.

Carafoli, F., Hussain, S.A., and Hohenester, E. (2012). Crystal structures of the network-forming short-arm tips of the laminin beta1 and gamma1 chains. PloS one 7, e42473.

Cheng, Y.S., Champliaud, M.F., Burgeson, R.E., Marinkovich, M.P., and Yurchenco, P.D. (1997). Self-assembly of laminin isoforms. The Journal of biological chemistry 272, 31525–31532.

Daniels, J.T., Geerling, G., Alexander, R.A., Murphy, G., Khaw, P.T., and Saarialho-Kere, U. (2003). Temporal and spatial expression of matrix metalloproteinases during wound healing of human corneal tissue. Experimental eye research 77, 653–664.

deHart, G.W., Healy, K.E., and Jones, J.C. (2003). The role of alpha3beta1 integrin in determining the supramolecular organization of laminin-5 in the extracellular matrix of keratinocytes. Experimental cell research 283, 67–79.

Dreier, B., Raghunathan, V.K., Russell, P., and Murphy, C.J. (2012). Focal adhesion kinase knockdown modulates the response of human corneal epithelial cells to topographic cues. Acta biomaterialia 8, 4285–4294.

Frank, D.E., and Carter, W.G. (2004). Laminin 5 deposition regulates keratinocyte polarization and persistent migration. Journal of cell science 117, 1351–1363.

Garbe, J.H., Gohring, W., Mann, K., Timpl, R., and Sasaki, T. (2002). Complete sequence, recombinant analysis and binding to laminins and sulphated ligands of the N-terminal domains of laminin alpha3B and alpha5 chains. Biochem J 362, 213–221.

Giard, D.J., Aaronson, S.A., Todaro, G.J., Arnstein, P., Kersey, J.H., Dosik, H., and Parks, W.P. (1973). In vitro cultivation of human tumors: establishment of cell lines derived from a series of solid tumors. Journal of the National Cancer Institute 51, 1417–1423.

Goldfinger, L.E., Hopkinson, S.B., deHart, G.W., Collawn, S., Couchman, J.R., and Jones, J.C. (1999). The alpha3 laminin subunit, alpha6beta4 and alpha3beta1 integrin coordinately regulate wound healing in cultured epithelial cells and in the skin. Journal of cell science 112 (Pt 16), 2615–2629.

Gonzales, M., Haan, K., Baker, S.E., Fitchmun, M., Todorov, I., Weitzman, S., and Jones, J.C. (1999). A cell signal pathway involving laminin-5, alpha3beta1 integrin, and mitogen-activated protein kinase can regulate epithelial cell proliferation. Molecular biology of the cell 10, 259–270.

Hamill, K.J., Hopkinson, S.B., DeBiase, P., and Jones, J.C. (2009a). BPAG1e maintains keratinocyte polarity through beta4 integrin-mediated modulation of Rac1 and cofilin activities. Molecular biology of the cell 20, 2954–2962.

Hamill, K.J., Hopkinson, S.B., Jonkman, M.F., and Jones, J.C. (2011). Type XVII collagen regulates lamellipod stability, cell motility, and signaling to Rac1 by targeting bullous pemphigoid antigen 1e to alpha6beta4 integrin. The Journal of biological chemistry 286, 26768–26780.

Hamill, K.J., Kligys, K., Hopkinson, S.B., and Jones, J.C. (2009b). Laminin deposition in the extracellular matrix: a complex picture emerges. Journal of cell science 122, 4409–4417.

Hamill, K.J., Langbein, L., Jones, J.C., and McLean, W.H. (2009c). Identification of a novel family of laminin N-terminal alternate splice isoforms: structural and functional characterization. The Journal of biological chemistry 284, 35588–35596.

Hampel, U., Garreis, F., Burgemeister, F., Essel, N., and Paulsen, F. (2018). Effect of intermittent shear stress on corneal epithelial cells using an in vitro flow culture model. Ocul Surf 16, 341–351.

Hashimoto, T., Amagai, M., Ebihara, T., Gamou, S., Shimizu, N., Tsubata, T., Hasegawa, A., Miki, K., and Nishikawa, T. (1993). Further analyses of epitopes for human monoclonal anti-basement membrane zone antibodies produced by stable human hybridoma cell lines constructed with Epstein-Barr virus transformants. The Journal of investigative dermatology 100, 310–315.

Hohenester, E., and Yurchenco, P.D. (2013). Laminins in basement membrane assembly. Cell adhesion & migration 7, 56–63.

Hopkinson, S.B., DeBiase, P.J., Kligys, K., Hamill, K., and Jones, J.C. (2008). Fluorescently tagged laminin subunits facilitate analyses of the properties, assembly and processing of laminins in live and fixed lung epithelial cells and keratinocytes. Matrix biology: journal of the International Society for Matrix Biology 27, 640–647.

Hopkinson, S.B., Findlay, K., deHart, G.W., and Jones, J.C. (1998). Interaction of BP180 (type XVII collagen) and alpha6 integrin is necessary for stabilization of hemidesmosome structure. The Journal of investigative dermatology 111, 1015–1022.

Hopkinson, S.B., Hamill, K.J., Wu, Y., Eisenberg, J.L., Hiroyasu, S., and Jones, J.C. (2014). Focal Contact and Hemidesmosomal Proteins in Keratinocyte Migration and Wound Repair. Adv Wound Care (New Rochelle) 3, 247–263.

Hopkinson, S.B., Riddelle, K.S., and Jones, J.C. (1992). Cytoplasmic domain of the 180-kD bullous pemphigoid antigen, a hemidesmosomal component: molecular and cell biologic characterization. The Journal of investigative dermatology 99, 264–270.

Hussain, S.A., Carafoli, F., and Hohenester, E. (2011). Determinants of laminin polymerization revealed by the structure of the alpha5 chain amino-terminal region. EMBO reports 12, 276–282.

Kabosova, A., Azar, D.T., Bannikov, G.A., Campbell, K.P., Durbeej, M., Ghohestani, R.F., Jones, J.C., Kenney, M.C., Koch, M., Ninomiya, Y., Patton, B.L., Paulsson, M., Sado, Y., Sage, E.H., Sasaki, T., Sorokin, L.M., Steiner-Champliaud, M.F., Sun, T.T., Sundarraj, N., Timpl, R., Virtanen, I., and Ljubimov, A.V. (2007). Compositional differences between infant and adult human corneal basement membranes. Investigative ophthalmology & visual science 48, 4989–4999.

Kaplan, N., Wang, J., Wray, B., Patel, P., Yang, W., Peng, H., and Lavker, R.M. (2019). Single-Cell RNA Transcriptome Helps Define the Limbal/Corneal Epithelial Stem/Early Transit Amplifying Cells and How Autophagy Affects This Population. Investigative ophthalmology & visual science 60, 3570–3583.

Kawamata, H., Nakashiro, K., Uchida, D., Harada, K., Yoshida, H., and Sato, M. (1997). Possible contribution of active MMP2 to lymph-node metastasis and secreted cathepsin L to bone invasion of newly established human oral-squamous-cancer cell lines. International journal of cancer. Journal international du cancer 70, 120–127.

Kivirikko, S., McGrath, J.A., Baudoin, C., Aberdam, D., Ciatti, S., Dunnill, M.G., McMillan, J.R., Eady, R.A., Ortonne, J.P., Meneguzzi, G., and et al. (1995). A homozygous nonsense mutation in the alpha 3 chain gene of laminin 5 (LAMA3) in lethal (Herlitz) junctional epidermolysis bullosa. Human molecular genetics 4, 959–962.

Koster, J., Geerts, D., Favre, B., Borradori, L., and Sonnenberg, A. (2003). Analysis of the interactions between BP180, BP230, plectin and the integrin alpha6beta4 important for hemidesmosome assembly. Journal of cell science 116, 387–399.

Langhofer, M., Hopkinson, S.B., and Jones, J.C. (1993). The matrix secreted by 804G cells contains laminin-related components that participate in hemidesmosome assembly in vitro. Journal of cell science 105 (Pt 3), 753–764.

Latvala, T., Tervo, K., and Tervo, T. (1995). Reassembly of the alpha 6 beta 4 integrin and laminin in rabbit corneal basement membrane after excimer laser surgery: a 12-month follow-up. The CLAO journal: official publication of the Contact Lens Association of Ophthalmologists, Inc 21, 125–129.

Ljubimov, A.V., Burgeson, R.E., Butkowski, R.J., Michael, A.F., Sun, T.T., and Kenney, M.C. (1995). Human corneal basement membrane heterogeneity: topographical differences in the expression of type IV collagen and laminin isoforms. Laboratory investigation; a journal of technical methods and pathology 72, 461–473.

Matsubara, M., Girard, M.T., Kublin, C.L., Cintron, C., and Fini, M.E. (1991a). Differential roles for two gelatinolytic enzymes of the matrix metalloproteinase family in the remodelling cornea. Dev Biol 147, 425–439.

Matsubara, M., Zieske, J.D., and Fini, M.E. (1991b). Mechanism of basement membrane dissolution preceding corneal ulceration. Investigative ophthalmology & visual science 32, 3221–3237.

Michopoulou, A., Montmasson, M., Garnier, C., Lambert, E., Dayan, G., and Rousselle, P. (2020). A novel mechanism in wound healing: Laminin 332 drives MMP9/14 activity by recruiting syndecan-1 and CD44. Matrix biology: journal of the International Society for Matrix Biology.

Momota, Y., Suzuki, N., Kasuya, Y., Kobayashi, T., Mizoguchi, M., Yokoyama, F., Nomizu, M., Shinkai, H., Iwasaki, T., and Utani, A. (2005). Laminin alpha3 LG4 module induces keratinocyte migration: involvement of matrix metalloproteinase-9. J Recept Signal Transduct Res 25, 1–17.

Nakano, A., Chao, S.C., Pulkkinen, L., Murrell, D., Bruckner-Tuderman, L., Pfendner, E., and Uitto, J. (2002). Laminin 5 mutations in junctional epidermolysis bullosa: molecular basis of Herlitz vs. non-Herlitz phenotypes. Human genetics 110, 41–51.

Nishiuchi, R., Takagi, J., Hayashi, M., Ido, H., Yagi, Y., Sanzen, N., Tsuji, T., Yamada, M., and Sekiguchi, K. (2006). Ligand-binding specificities of laminin-binding integrins: a comprehensive survey of laminin-integrin interactions using recombinant alpha3beta1, alpha6beta1, alpha7beta1 and alpha6beta4 integrins. Matrix biology: journal of the International Society for Matrix Biology 25, 189–197.

Odenthal, U., Haehn, S., Tunggal, P., Merkl, B., Schomburg, D., Frie, C., Paulsson, M., and Smyth, N. (2004). Molecular analysis of laminin N-terminal domains mediating self-interactions. The Journal of biological chemistry 279, 44504–44512.

Ofuji, K. (2000). Differential tyrosine phosphorylation of paxillin in human corneal epithelial cells on extracellular matrix proteins. Japanese journal of ophthalmology 44, 189.

Pal-Ghosh, S., Blanco, T., Tadvalkar, G., Pajoohesh-Ganji, A., Parthasarathy, A., Zieske, J.D., and Stepp, M.A. (2011a). MMP9 cleavage of the beta4 integrin ectodomain leads to recurrent epithelial erosions in mice. Journal of cell science 124, 2666–2675.

Pal-Ghosh, S., Pajoohesh-Ganji, A., Tadvalkar, G., Kyne, B.M., Guo, X., Zieske, J.D., and Stepp, M.A. (2016). Topical Mitomycin-C enhances subbasal nerve regeneration and reduces erosion frequency in the debridement wounded mouse cornea. Experimental eye research 146, 361–369.

Pal-Ghosh, S., Pajoohesh-Ganji, A., Tadvalkar, G., and Stepp, M.A. (2011b). Removal of the basement membrane enhances corneal wound healing. Experimental eye research 93, 927–936.

Pal-Ghosh, S., Tadvalkar, G., Jurjus, R.A., Zieske, J.D., and Stepp, M.A. (2008). BALB/c and C57BL6 mouse strains vary in their ability to heal corneal epithelial debridement wounds. Experimental eye research 87, 478–486.

Pfendner, E.G., and Lucky, A.W. (1993). Junctional Epidermolysis Bullosa. In: GeneReviews((R)), eds. M.P. Adam, H.H. Ardinger, R.A. Pagon, S.E. Wallace, L.J.H. Bean, K. Stephens, and A. Amemiya, Seattle (WA).

Pullar, C.E., Baier, B.S., Kariya, Y., Russell, A.J., Horst, B.A., Marinkovich, M.P., and Isseroff, R.R. (2006). beta4 integrin and epidermal growth factor coordinately regulate electric field-mediated directional migration via Rac1. Molecular biology of the cell 17, 4925–4935.

Purvis, A., and Hohenester, E. (2012). Laminin network formation studied by reconstitution of ternary nodes in solution. The Journal of biological chemistry 287, 44270–44277.

Reuten, R., Patel, T.R., McDougall, M., Rama, N., Nikodemus, D., Gibert, B., Delcros, J.G., Prein, C., Meier, M., Metzger, S., Zhou, Z., Kaltenberg, J., McKee, K.K., Bald, T., Tuting, T., Zigrino, P., Djonov, V., Bloch, W., Clausen-Schaumann, H., Poschl, E., Yurchenco, P.D., Ehrbar, M., Mehlen, P., Stetefeld, J., and Koch, M. (2016). Structural decoding of netrin-4 reveals a regulatory function towards mature basement membranes. Nature communications 7, 13515.

Robertson, D.M., Li, L., Fisher, S., Pearce, V.P., Shay, J.W., Wright, W.E., Cavanagh, H.D., and Jester, J.V. (2005). Characterization of growth and differentiation in a telomerase-immortalized human corneal epithelial cell line. Invest. Ophthalmol. Vis. Sci. 46, 470–478.

Schittny, J.C., and Yurchenco, P.D. (1990). Terminal short arm domains of basement membrane laminin are critical for its self-assembly. The Journal of cell biology 110, 825–832.

Schlotzer-Schrehardt, U., Dietrich, T., Saito, K., Sorokin, L., Sasaki, T., Paulsson, M., and Kruse, F.E. (2007). Characterization of extracellular matrix components in the limbal epithelial stem cell compartment. Experimental eye research 85, 845–860.

Schneiders, F.I., Maertens, B., Bose, K., Li, Y., Brunken, W.J., Paulsson, M., Smyth, N., and Koch, M. (2007). Binding of netrin-4 to laminin short arms regulates basement membrane assembly. The Journal of biological chemistry 282, 23750–23758.

Sehgal, B.U., DeBiase, P.J., Matzno, S., Chew, T.L., Claiborne, J.N., Hopkinson, S.B., Russell, A., Marinkovich, M.P., and Jones, J.C. (2006). Integrin beta4 regulates migratory behavior of keratinocytes by determining laminin-332 organization. The Journal of biological chemistry 281, 35487–35498.

Staquicini, F.I., Dias-Neto, E., Li, J., Snyder, E.Y., Sidman, R.L., Pasqualini, R., and Arap, W. (2009). Discovery of a functional protein complex of netrin-4, laminin gamma1 chain, and integrin alpha6beta1 in mouse neural stem cells. Proc Natl Acad Sci U S A 106, 2903–2908.

Suzuki, K., Tanaka, T., Enoki, M., and Nishida, T. (2000). Coordinated reassembly of the basement membrane and junctional proteins during corneal epithelial wound healing. Investigative ophthalmology & visual science 41, 2495–2500.

Taniguchi, Y., Ido, H., Sanzen, N., Hayashi, M., Sato-Nishiuchi, R., Futaki, S., and Sekiguchi, K. (2009). The C-terminal region of laminin beta chains modulates the integrin binding affinities of laminins. The Journal of biological chemistry 284, 7820–7831.

Taniguchi, Y., Takizawa, M., Li, S., and Sekiguchi, K. (2020). Bipartite mechanism for laminin-integrin interactions: Identification of the integrin-binding site in LG domains of the laminin alpha chain. Matrix biology: journal of the International Society for Matrix Biology 87, 66–76.

Troughton, L.D., Reuten, R., Sugden, C.J., and Hamill, K.J. (2020a). Laminin N-terminus α31 protein distribution in adult human tissues. bioRxiv, 2020.2005.2021.108134.

Troughton, L.D., Zech, T., and Hamill, K.J. (2020b). Laminin N-terminus α31 is upregulated in invasive ductal breast cancer and changes the mode of tumour invasion. bioRxiv, 2020.2005.2028.120964.

Tsuruta, D., Hashimoto, T., Hamill, K.J., and Jones, J.C. (2011). Hemidesmosomes and focal contact proteins: functions and cross-talk in keratinocytes, bullous diseases and wound healing. Journal of dermatological science 62, 1–7.

Turner, C.E. (2000). Paxillin and focal adhesion signalling. Nature cell biology 2, E231–236.

Walko, G., Castanon, M.J., and Wiche, G. (2015). Molecular architecture and function of the hemidesmosome. Cell and tissue research 360, 363–378.

Wang, W., Zuidema, A., Te Molder, L., Nahidiazar, L., Hoekman, L., Schmidt, T., Coppola, S., and Sonnenberg, A. (2020). Hemidesmosomes modulate force generation via focal adhesions. The Journal of cell biology 219.

Yamamoto, N., Yamamoto, N., Petroll, M.W., Cavanagh, H.D., and Jester, J.V. (2005). Internalization of Pseudomonas aeruginosa is mediated by lipid rafts in contact lens-wearing rabbit and cultured human corneal epithelial cells. Investigative ophthalmology & visual science 46, 1348–1355.

Yanez-Soto, B., Leonard, B.C., Raghunathan, V.K., Abbott, N.L., and Murphy, C.J. (2015). Effect of Stratification on Surface Properties of Corneal Epithelial Cells. Investigative ophthalmology & visual science 56, 8340–8348.

Yurchenco, P.D., and Cheng, Y.S. (1993). Self-assembly and calcium-binding sites in laminin. A three-arm interaction model. The Journal of biological chemistry 268, 17286–17299.

Yurchenco, P.D., Tsilibary, E.C., Charonis, A.S., and Furthmayr, H. (1985). Laminin polymerization in vitro. Evidence for a two-step assembly with domain specificity. The Journal of biological chemistry 260, 7636–7644.

