## Supplemental Figure legends for "Laminin N-terminus α31 regulates keratinocyte adhesion and migration through modifying the organisation and proteolytic processing of laminin 332"

### Supplemental material

**Supplemental Figure 1: Cell line validation. (**A) hTCEpi corneal epithelial cells, human retinal pericytes (HRP) or human dermal fibroblasts were plated overnight on glass coverslips then fixed and processed with antibodies against keratin 12 and vimentin. Scale bar 20 µm. (B) DNA was extracted from HaCaT, BHY, and A549 cells and processed for short tandem repeat cell fingerprinting.

**Supplemental Figure 2: Epidermal and oral keratinocyte gap closure is slowed in cells transduced with LaNt α31.** HaCat epidermal keratinocytes (A and B) or BHY oral keratinocytes (C and D) were induced to express eGFP (+eGFP) or LaNt α31-eGFP (+LaNt α31) then plated overnight on plastic dishes in Ibidi® 2-well culture inserts, which were then removed at the beginning of the assay. (A and C) Representative images at 0 h and 16 h, with yellow lines indicating cell sheet margins. Scale bar 100 µm. (B and D) Quantification of wound area remaining at 16 h as percentage of initial gap. Each point represents an independent experimental repeat with 2 or 3 technical repeats per assay with a line indicating the mean. * denotes p<0.05 compared with both controls as determined by one-way ANOVA with Bonferroni post-hoc test.

**Supplemental Figure 3: Laminin isoform transcript expression comparison.** Total RNA was extracted from hTCEpi, HaCaT or BHY cells and RT-qPCR performed with primers specific to each laminin encoding gene. Bar chart represents the abundance of each transcript relative to LAMA3A levels within each cell line.

**Supplemental Figure 4 Induced LaNt α31 expression changes LM332 organisation.** Primary corneal epithelial cells (pCEC, A, B) or hTCEpi (C, D and E) were transduced with eGFP (eGFP) or LaNt α31-eGFP (+LaNt α31) plated overnight on glass coverslips then fixed and processed for indirect immunofluorescence microscopy with antibodies against laminin α3 (LMα3), laminin β3 (LMβ3), laminin γ2 (LMγ2), laminin α5 (LMα5), or polyclonal antibodies raised against LM332. The lung adenocarcinoma line A549 are included as a control for LMα5 deposition. Scale bars 20 µm. (B) Cluster area of LMα3 in pixels measured from four primary cell donors with a minimum of 10 cells per donor. Boxes in represent 25th and 75th percentile, whiskers 5th and 95th percentile. Red * denote P<0.05 for +LaNt α31 compared with both the +GFP and non-transduced pCEC populations as determined by one way ANOVA followed by Tukey's post hoc test.

**Supplemental Figure 5 LaNt α31 is organised into discrete patches at the bottom of cells.** (A) hTCEpi cells transduced with LaNt α31-eGFP adenovirus were plated for 16 h on glass-bottomed dishes then imaged using epifluorescence (left) and total internal reflection fluorescence (TIRF) microscopy (right). (B) LaNt α31-eGFP expressing hTCEpi were plated onto glass coverslips and 16h later the cellular material removed through ammonium hydroxide treatment then remaining ECM material imaged. Phase contrast images in the inset in (B) show the absense of cells. Scale bars 10 µm.

**Supplemental Figure 6 LMβ3-mCherry co localises with LMα3 and LMγ2** hTEpi cells were transduced with LMβ3-mCherry adenovirus, plated overnight onto glass coverslips then processed for indirect immunofluorescence with antibodies against LMα3, LMβ3, LMγ2 or isotype IgG control (left panels). Right panels are red fluorescent signal from mCherry signal. Scale bar 10 µm.

**Supplemental Figure 7 Integrin β4 co-distributes with LM332.** Non-transduced, +eGFP, or +LaNt α31 hTCEpi were plated overnight on glass coverslips then processed for indirect immunofluorescence with polyclonal antibodies against LM332 or monoclonal antibodies against integrin β4. Scale bar 20 µm.

**Supplemental Figure 8 Inhibiting matrix metalloproteinases rescues the LaNt α31 expression effects on hemidesmosome formation and focal adhesion distribution** +LaNt α31 hTCEpi cells were plated overnight with broad spectrum MMP inhibitor (MMPi) or serine protease inhibitor (non-MMPi) and coverslips processed for indirect immunofluorescence with antibodies against bulous pemphigoid antigen 1e (BPAG1e) and β4 integrin (A) or paxillin and β4 integrin (B). Scale bars 20 µm.
