## Supplemental Methods for "Laminin N-terminus α31 regulates keratinocyte adhesion and migration through modifying the organisation and proteolytic processing of laminin 332"

### DNA extraction for cell line authentication

### For short tandem repeat (STR) loci-cell line authentication, cells were seeded at 2.5 x 10^5^/ well of a 6-well plate (Greiner-BioOne) and DNA extracted the following day using the Monarch® Genomic DNA Purification Kit from New England BioLabs (NEB inc, Ipswich, Massachusetts, USA). DNA was then sent to the cell line authentication service for cell line validation at Liverpool Bio-Innovation Hub, University of Liverpool.

### RT-qPCR

### For LM isoform-specific RT-qPCR cells were seeded at 2.5 x 10^5^/ well of a 6-well plate and RNA extracted after 16 h using RNeasy mini-prep spin columns from Qiagen (Qiagen, Venlo, Netherlands). RNA quantities and purities were measured using a Nanodrop 2000™ (ThermoFisher) accepting an optical density 260/280 ratio of between 1.9 and 2.1. 1 μg of total RNA was reverse transcribed using Precision nanoScript™ 2 Reverse Transcriptase (Primerdesign, Camberley, UK) using random hexamers and oligo-DT primers under the following conditions: primer annealing at 65°C for 5 min followed by transcriptase extension at 25°C for 5 min and 42°C for 20 min, before inactivating the enzyme at 75°C for 10 min. 5 ng of cDNA was used for each 10 μL qRT-PCR reaction containing 1 μM primer pair (Supplemetal Table 1) and 5 μL Precision Plus qPCR Mastermix (Primerdesign). The following run protocol was used for all qRT-PCR reactions: 1 cycle of 95°C for 2 min, 40 cycles of 95°C for 15 secs followed by 1 min at 60°C, with a final melt curve analysis. All qRT-PCR was performed using Roche Lightcyler 480™ or Lightcycler 96™ (Roche, Basel, Switzerland).
