## Supplemental Table 1 for "Laminin N-terminus α31 regulates keratinocyte adhesion and migration through modifying the organisation and proteolytic processing of laminin 332"

Primer information for **laminin and reference transcripts**

| **Isoform** | **Direction** | **Sequence** | **Location** | **Amplicon Size** |
| --- | --- | --- | --- | --- |
| LAMA1 | Forward  Reverse | 5'- GGCTCTGTGACTGCAAACCAAACGTG– 3'  5'- CAGCCATGGCCTGAGTCCAGC– 3' | Exon 20-21 | 88 bp |
| LAMA2 | Forward  Reverse | 5'- CCTCTTGTGTCGCAGAAGGACTTGACG– 3'  5'- GCCTGGACTGCCAGTATAGCCAG– 3' | Exon 31-32 | 114 bp |
| LAMA3A | Forward  Reverse | 5'- GCCTCCCGGTCAAAGTCAACTGC– 3'  5'- GCAGGGAACACACCGTCCGGTATAC– 3' | Exon 39-40 | 118 bp |
| LAMA3B | Forward  Reverse | 5'- GCCACCTGTGTCTCCTTGGCC– 3'  5'- CCAGGTGTGGTACACGTCCTCTCA– 3' | Exon 27-28 | 181 bp |
| LAMA4 | Forward  Reverse | 5'- CCAGACTCAGTGATGCCGTTAAGCAAC– 3'  5'- GGCTTCCTCGGTGATCAGTCTAGACTGC– 3' | Exon 17-18 | 95 bp |
| LAMA5 | Forward  Reverse | 5'- AGCGGTGTGACGTGTGTGCC– 3'  5'- CTGCCGCGCTGCAGTCACAAT– 3' | Exon 10-12 | 132 bp |
| LAMB1 | Forward  Reverse | 5'- CATGAGACCCTGAATCCTGACAGCC– 3'  5'- GCATAGCAGCTGGACGGAATGTCTTGAAAGTC– 3' | Exon 4-6 | 190 bp |
| LAMB2 | Forward  Reverse | 5'- GCCTGTGTGGGCATTTGGTGC– 3'  5'- CCACTTCAGATGCAGCTTGTAGGAGA – 3' | Exon 15-16 | 132 bp |
| LAMB3 | Forward  Reverse | 5'- GGAGAAAGAACGGCAGAACACACAGC– 3'  5'- GGCAGGGCAAAACACAAGAGGAAGAA– 3' | Exon 1-3 | 91 bp |
| LAMC1 | Forward  Reverse | 5'- CCCAGCTCCATCAACCTCACGC– 3'  5'- CCGTGTGCGCTTGTAAATGGCAAAGC– 3' | Exon 1-2 | 141 bp |
| LAMC2 | Forward  Reverse | 5'- CTGCCTCTGCTTCTCGCTCCTC– 3'  5'- CTGCCAGGAGTTCCCATTGCTTAGACAGA– 3' | Exon 1-2 | 85 bp |
| LAMC3 | Forward  Reverse | 5'- CCAGCATGGCACCTGTGACC – 3'  5'- CCATAGAAACCTGGCAAACAGCGTTC – 3' | Exon 12-13 | 96 bp |
