## Supplemental Movie Legends for "Laminin N-terminus α31 regulates keratinocyte adhesion and migration through modifying the organisation and proteolytic processing of laminin 332"

### Supplemental Movie 1

hTCEpi co-transduced with LaNt α31-eGFP and LMβ3-mCherry were plated overnight on uncoated glass-bottomed dishes and imaged by confocal microscopy every 20 min. Left panel LaNt α31-eGFP, middle panel LMβ3-mCherry with signals inverted, right panel, merged LaNt α31-eGFP (pseudocoloured green) and LMβ3 mCherry (pseudocoloured magenta).

### Supplemental Movie 2

hTCEpi transduced with LaNt α31-eGFP and LMβ3-mCherry were plated overnight in a plastic cloning ring on an uncoated glass-bottomed dish and imaged by confocal microscopy every 10 min for 16 h. Left panel LaNt α31-eGFP, middle panel LMβ3-mCherry with signals inverted, right panel, merged LaNt α31-eGFP (pseudocoloured green) and LMβ3-mCherry (pseudocoloured magenta).
